# The small GTPase Arf1 regulates ATP synthesis and mitochondria homeostasis by modulating fatty acid metabolism

**DOI:** 10.1101/2022.01.26.477847

**Authors:** Ludovic Enkler, Mirjam Pennauer, Viktoria Szentgyörgyi, Cristina Prescianotto-Baschong, Isabelle Riezman, Aneta Wiesyk, Roza Kucharczyk, Martin Spiess, Howard Riezman, Anne Spang

## Abstract

Lipid mobilization through fatty acid β-oxidation is a central process essential for energy production during nutrient shortage. In yeast, this catabolic process starts in the peroxisome from where β-oxidation products enter mitochondria and fuel the TCA cycle. Little is known about the physical and metabolic cooperation between these organelles. We found that expression of fatty acid transporters and of the rate-limiting enzyme involved in β-oxidation are decreased in cells expressing a hyperactive mutant of the small GTPase Arf1, leading to an accumulation of fatty acids in lipid droplets. As a consequence, mitochondria became fragmented and ATP synthesis decreased. Genetic and pharmacological depletion of fatty acids phenocopied the *arf1* mutant mitochondrial phenotype. Although β-oxidation occurs mainly in mitochondria in mammals, Arf1’s role in fatty acid metabolism is conserved. Together, our results indicate that Arf1 integrates metabolism into energy production by regulating fatty acid storage and utilization, and presumably organelle contact-sites.

## Introduction

Intracellular compartmentalization of metabolic processes involves deep and well-orchestrated inter-organelle communications to coordinate cellular functions. This requires homeostatic control of lipid, ion, and metabolite transfer between organelles, as well as between organelles and the plasma membrane [1-6]. Exchanges are established by two means, namely vesicular transport for organelles in the secretory pathway by means of kissing and fusing [1], or by an elaborate network of membranes dedicated to establish membrane contact sites [5, 7].

Mitochondria form contacts with almost every organelle in the cell [2, 3, 8]. They establish functional interactions with peroxisomes [9] and with lipid droplets (LDs; [10, 11]) to ensure fatty acid (FA) metabolism and ATP production. Both in yeast and mammals, lipids are stored in LDs in the form of triacylglycerol (TAG) and sterol esters (SE). Under nutrient shortage, FAs are released from LDs by lipolysis and metabolized by β-oxidation solely in peroxisome in yeast, or both in peroxisomes and mitochondria in mammalian cells. Subsequently, shortened acyl-CoA (or acetylcarnitine) is transferred from peroxisomes to mitochondria by an unknown mechanism [9, 12], where it will be used to fuel the TCA cycle and the respiratory chain (RC) complexes for oxidative phosphorylation (OXPHOS). Hence, LDs stay in close proximity to peroxisomes and mitochondria in yeast and mammals for efficient transfer of metabolites [13-17]. Increasing evidence points towards the fact that perturbed contact sites between these organelles and mitochondria are important parts of metabolic syndromes, liver disease, and cancers highlighting their central role in cellular homeostasis [8, 18, 19]. Nevertheless, there is an enormous lack of mechanistic insights on how contact sites are organized and which proteins are involved in metabolite transfer allowing proper lipid flux between organelles to insure effective energy synthesis.

Arf1 is a master regulator of vesicle formation (i.e COPI- and clathrin-coated vesicles) at the Golgi [20, 21] and its activity is modulated by ArfGAPs (GTPase activating proteins) and ArfGEFs (guanine nucleotide exchange factors) stimulating GTP hydrolysis or GDP-to-GTP exchange, respectively. Over the past few years, additional functions of Arf1 have been identified. We and others have shown that Arf1 regulates mRNA trafficking [22, 23], mTORC1 activity [24, 25], and mitochondrial dynamics and transport [26-28]. However, it remains still unclear how Arf1 specifically regulates mitochondrial dynamics. While we and others have shown that knocking-down *ARF1* in *C. elegans* or knocking out ARF1 in Hela cells leads to mitochondrial hyperconnectivity [26, 28], we found that mitochondria were fragmented and globular in the yeast *arf1-11* mutant [26, 29]. Therefore, these opposing results led us to surmise that Arf1 might have roles at mitochondria beyond the regulation of their dynamics.

The Arf1/COPI machinery has also been implicated in lipid metabolism by governing lipolysis, LD morphology, protein recruitment, phospholipid removal, and the formation of ER-LD bridges [30-36]. Additionally, some reports provided evidence that active GTP-bound Arf1 and the COPI machinery are recruited onto peroxisomes [37, 38], and suggested that Arf1 is involved in peroxisome proliferation [38-40]. Moreover, Arf1 localization appears to depend on the peroxisomal protein Pex35 and potentially Pex11 [37, 40]. However, Arf1 function on peroxisomes and how this contributes to the metabolism of FAs remains still elusive.

Here, we show that Arf1 couples FA β-oxidation to mitochondrial ATP synthesis. Using a predominantly active mutant of Arf1, *arf1-11*, we demonstrate that Arf1 activity regulates expression of long-chain fatty acid transporters Pxa1/Pxa2 and of Pox1, the first and rate-limiting enzyme involved in β-oxidation in yeast. As a consequence, *arf1-11* leads to an increased level of TAG in LDs, and to reduced lipid transfer from peroxisomes in yeast, or from LDs in mammalian cells, to mitochondria. This conserved mechanism is essential to sustain endomembrane homeostasis and ATP synthesis by mitochondria. In addition, our work suggests that Arf1 activity drives both mitochondrial fusion and fission in yeast, thereby explaining the different findings on the role of Arf1 in mitochondria dynamics [26, 28]. Our data provide evidence for at least two distinct roles of Arf1 on mitochondria: one in the regulation of mitochondrial dynamics and one in the transfer of FA to mitochondria.

## Results

### Arf1 regulates mitochondrial fusion and fission

To better understand the discrepancies between the hyperconnectivity of mitochondria observed in mammalian cells and *C. elegans* [26, 28] and the globular, fragmented mitochondria in the yeast *arf1-11* mutant strain, we aimed to measure mitochondrial fission and fusion activity in the *ARF1* and *arf1-11* strains. For clarity, we will use the prefix ‘y’ for all yeast and ‘m’ for all mammalian genes and proteins. yArf1-11 bears mutations in the GTP binding domain (K38T and E132D) and in the C-terminal tail (L173S) (**Figure 1A**, [29]). The mutant grows similar to y*ARF*1 at the permissive temperature (23°C) but is lethal when grown at the restrictive temperature (37°C) (**Figure 1B**). To determine the optimal condition for imaging, we shifted y*arf1-11* cells for various times to the restrictive temperature and determined their growth and morphology over time. The growth of y*arf1-11* cells slowed down already shortly after shift to the restrictive temperature, and the culture never reached the exponential growth phase (**Figure 1C**). Nevertheless, the morphology of y*arf1-11* cells appeared normal for up to 1 hr at 37°C (**Figure S1A**). Based on this analysis, we decided to shift cells for 30 min to 37°C prior to imaging.

**Figure 1.**
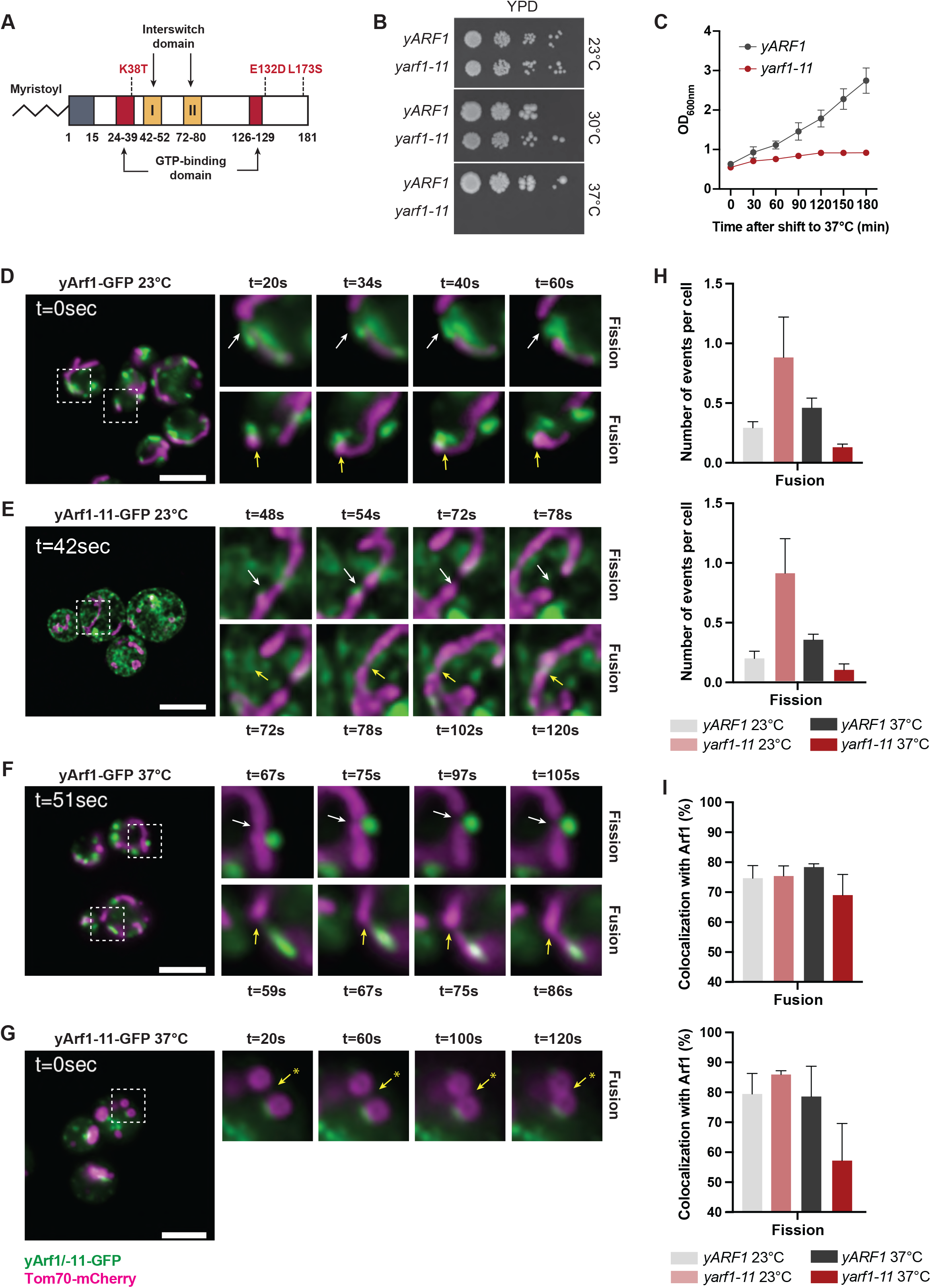
yArf1 regulates mitochondria fusion and fission. **A)** Schematic of the thermo-sensitive mutant Arf1-11 in yeast (Yahara et al., 2001). Amino acid coordinates are indicated in bold below the protein, and corresponding mutated amino acids in red. **B)** Growth test of y*ARF1* and y*arf1-11* strains on rich media (YPD) and incubated at 23°C, 30°C and 37°C. **C)** Cell viability assay of y*ARF1* and y*arf1-11* strains performed after shifting cells to 37°C. Optical densities (OD) were measured at regular time points. **D-G)** Single time-point images of movies done with strains expressing yArf1-GFP or yArf1-11-GFP together with the mitochondrial protein Tom70 fused to mCherry at 23°C or 37°C. White arrows indicate sites of fission and yellow arrows fusion. * indicates a fusion event independent of Arf1 in **G**. Scale bar 5 µm. **H-I)** Measurements of mitochondrial fusion and fission events per cell (**H**) and the frequency of events where yArf1 is involved (**I**). Mean and standard deviation are shown. See also Figure S1

We visualized mitochondria by genomically tagging Tom70 with mCherry in strains expressing genomically tagged yArf1-GFP or yArf1-11-GFP (**Table S1**). In yArf1-GFP cells, we could readily detect mitochondrial fission and fusion events at both 23°C and 37°C (**Figure 1D, F and H; Movie 1 and 2**). Importantly, we detected yArf1 at the sites on mitochondria where either fission or fusion occurred. (**Figure 1D, F and I, Figure S2A, C**). To our surprise, in *yarf1-11* cells, the number of fission and fusion events was higher than in yArf1 cells at the permissive temperature (23°C) (**Figure 1E, H, Figure S2B; Movie 3**). In contrast at the restrictive temperature (37°C), mitochondrial dynamics was greatly reduced (**Figure 1G, H, Figure S2D; Movie 4**). However, yArf1-11-GFP was still mostly present at the remaining fusion sites and to a somewhat lesser extent at the fission sites under these conditions (**Figure 1E, G, I**). Taken together our data suggest that yArf1 is required for both mitochondrial fusion and fission reconciling the findings in mammalian cells, *C. elegans*, and yeast.

### yArf1-11 is a hyperactive mutant that localizes to the ER and lipid droplets

However, we were puzzled by the difference in mitochondrial morphology in these experimental systems. A main difference between the experiments in yeast, *C. elegans*, and mammalian cells was that in yeast we used a mutant, while in the other systems knockdowns or knockouts were performed [26, 28]. yArf1 is a small GTPase that cycles between an active GTP-bound state and an inactive GDP-bound state. Since two of the three mutations of y*arf1-11* are located within or in close proximity to the GTP-binding domain, we explored the possibility that the y*arf1-11* allele could be linked to Arf1 activity. To test this hypothesis, we asked whether GTP binding was impaired in yArf1-11. To this end, we expressed and purified the GAT domain of the Arf effector Gga2 (Gga2^GAT^) known to specifically recruit yArf1 in its GTP-bound form (**Figure S1H**) [41]. Soluble (S100) and pellet (P100) fractions from yArf1-GFP and yArf1-11-GFP cells grown at 23°C or 37°C were incubated with Gga2^GAT^ and Arf1 binding was assessed. The yArf1-11 mutant protein in the P100 fraction was more efficiently retained by Gga2^GAT^ compared to yArf1 regardless of the temperature (**Figure 2A**). At 23°C, yArf1-11 in the S100 fraction also bound Gga2^GAT^ although to a lower extent. To confirm that retention was not induced by unspecific binding due to the GFP tag, we performed similar experiments using cells with untagged yArf1 and yArf1-11. Again, membrane bound yArf1-11 was more prominently retained by Gga2^GAT^ than yArf1, independent of the temperature (**Figure 2B**). Under these conditions, untagged soluble yArf1-11 in the S100 fraction was retained by Gga2^GAT^ similarly to what we observed with the tagged yArf1 proteins (**Figure 2A**). Our results indicate that yArf1-11 is mostly in the active conformation already at 23°C, that GTP-binding does not change upon shift to the restrictive temperature, and hence that y*arf1-11* is a gain-of-function mutant. This result finally explains the difference between the experimental systems. Loss of Arf1 function yields hyperfused mitochondria, while a gain-of-function mutation results in globular mitochondria.

**Figure 2.**
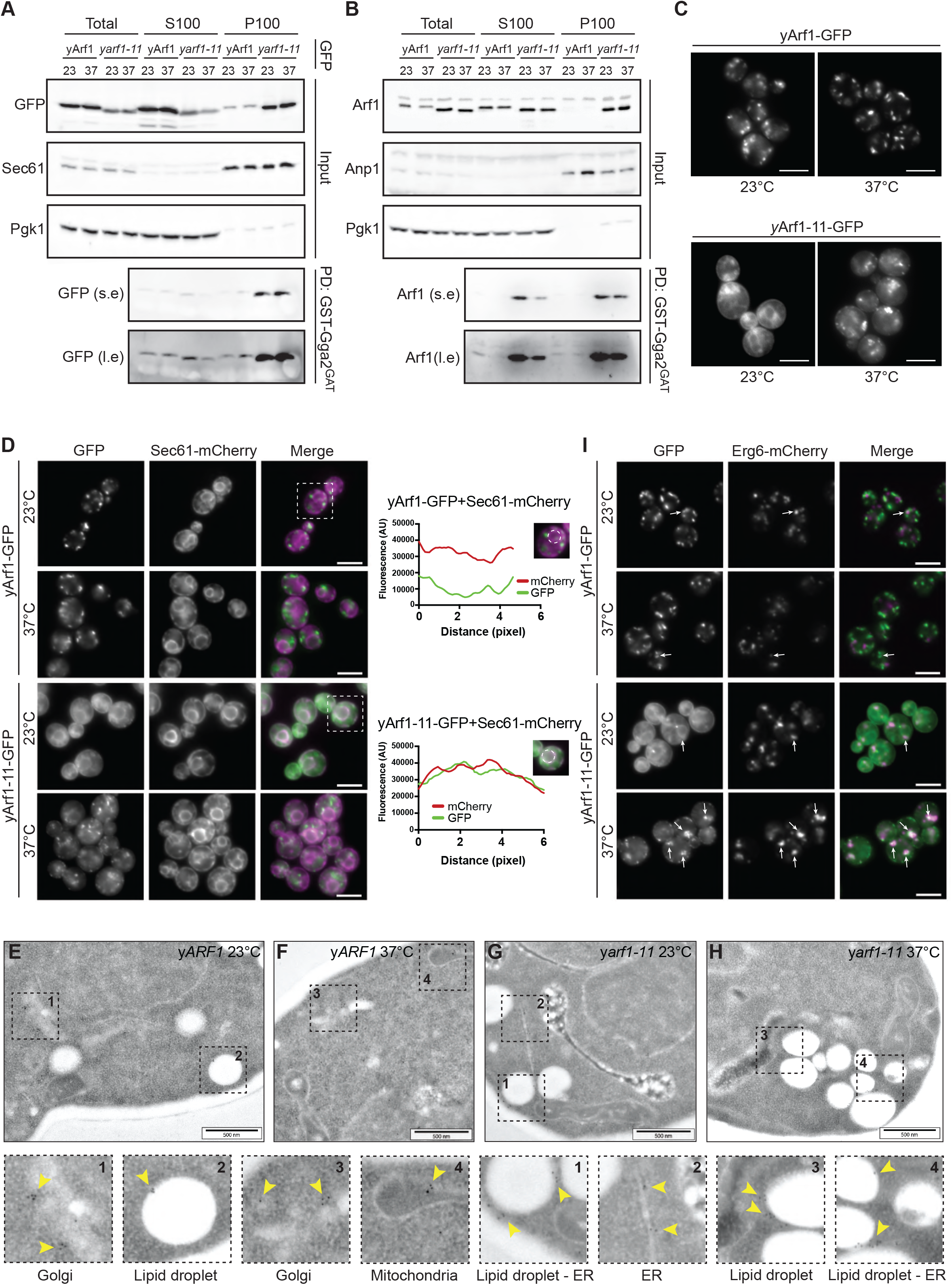
yArf1-11 is a hyperactive mutant that localizes to the ER and lipid droplets. **A-B)** Active yArf1 pull-down and detection experiments done with strains expressing yArf1 and yArf1-11 fused to GFP (**A**) or endogenous untagged yArf1 and yArf1-11 (**B**). Protein extracts from soluble (S100) or pellet (P100) fractions from y*ARF1* and y*arf1-11* cells grown at 23°C or 37°C were incubated with equal amount of purified GST-tagged GAT domain of Gga2 (Gga2^GAT^). Sec61 and Anp1 were used as ER marker and Pgk1 as cytosolic marker. s.e: short exposure; l.e: long exposure; PD: pull-down. **C)** Localization of wild type yArf1 and yArf1-11 C-terminally fused to GFP in the yeast *Saccharomyces cerevisiae*. Cells were incubated either at 23°C or shifted at 37°C for 30 min. **D)** Co-localization of yArf1-GFP and yArf1-11-GFP with the ER marker Sec61 tagged with mCherry grown at 23°C and 37°C. Cells highlighted by dotted squares were used to look for GFP and mCherry co-localization. Fluorescence intensities of each channel were measured on a circle drawn around the perinuclear ER and are shown here as arbitrary units (AU). **E-H)** Transmission electron microscopy of y*ARF1* and y*arf1-11* strains grown either at 23°C or shifted at 37°C for 30 min. yArf1 and yArf1-11 localizations were highlighted by immunogold labelling, and dotted squares shows enlargements of specific Arf1 localizations. Scale bars: 500 nm. **I)** Co-localization of yArf1-GFP and yArf1-11-GFP with the LD marker Erg6 tagged with mCherry grown at 23°C and 37°C. Arrows indicate sites of co-localization between the yArf1/yArf1-11 and LD. Scale bar 5 µm. See also Figure S2

When we recorded the movies on mitochondrial dynamics, we noticed that the yArf1-11 localization pattern was different to that of yArf1. This could either be due to a difference in Golgi morphology in y*arf1-11*, where the bulk of yArf1 is localized, or yArf1-11 might localize to different organelles. yArf1-GFP mainly localized, as expected, to the *cis*- and *trans*-Golgi (**Figure S1B, D**), at 23°C. yArf1-11-GFP was surprisingly present at the cell periphery and around the nucleus, conspicuously similar to endoplasmic reticulum localization, and to a much lesser extent at the *cis*- and *trans*-Golgi compartments (**Figure 2C, E**). We confirmed the ER localization with the ER marker Sec61-mCherry (**Figure 2D**) and by immuno electron microscopy (immuno-EM) with antibodies against Arf1 (**Figure 2E-H**). Since y*arf1-11* mutant cells do not have a growth defect at 23°C (**Figure 1B**), Arf1-11’s ER localization does not seem to be detrimental. When we shifted the cells to 37°C, however, yArf1-11 was massively relocated to puncta, which did not correspond to Golgi compartments (**Figure 2C, Figure S1C, E**). Arf1 has been reported to be localized also to LDs and mitochondria [30, 31, 33, 36, 42, 43], which we also observed irrespective of growth temperature (**Figure 2E and F**). Therefore, we tested whether these puncta corresponded to LDs by co-staining with the LD marker Erg6-mCherry and by immuno-EM, which was indeed the case (**Figure 2I, Figure S2F, G**). Of note, the number of LDs appeared to be increased and clustered in y*arf1-11* compared to wild-type (**Figure 2G, H, I, Figure S1G**). From these observations, we conclude that the bulk of yArf1-11 mainly localizes to the ER at the permissive temperature and to LDs at the restrictive temperature.

### yArf1-11 localization on LDs induces mitochondrial fragmentation

Since yArf1-11 is a gain-of-function mutant, we hypothesized that the dominant active version of Arf1, yArf1Q71L might localize in a similar fashion. Indeed, yArf1Q71L localized in a fashion reminiscent of yArf1-11 at the ER and in smaller puncta that were quite distinct from the Golgi localization observed with yArf1 and the dominant inactive version yArf1T31N (**Figure 3A**). Therefore, the active form of yArf1 can be found on the ER and most likely also on LDs.

**Figure 3.**
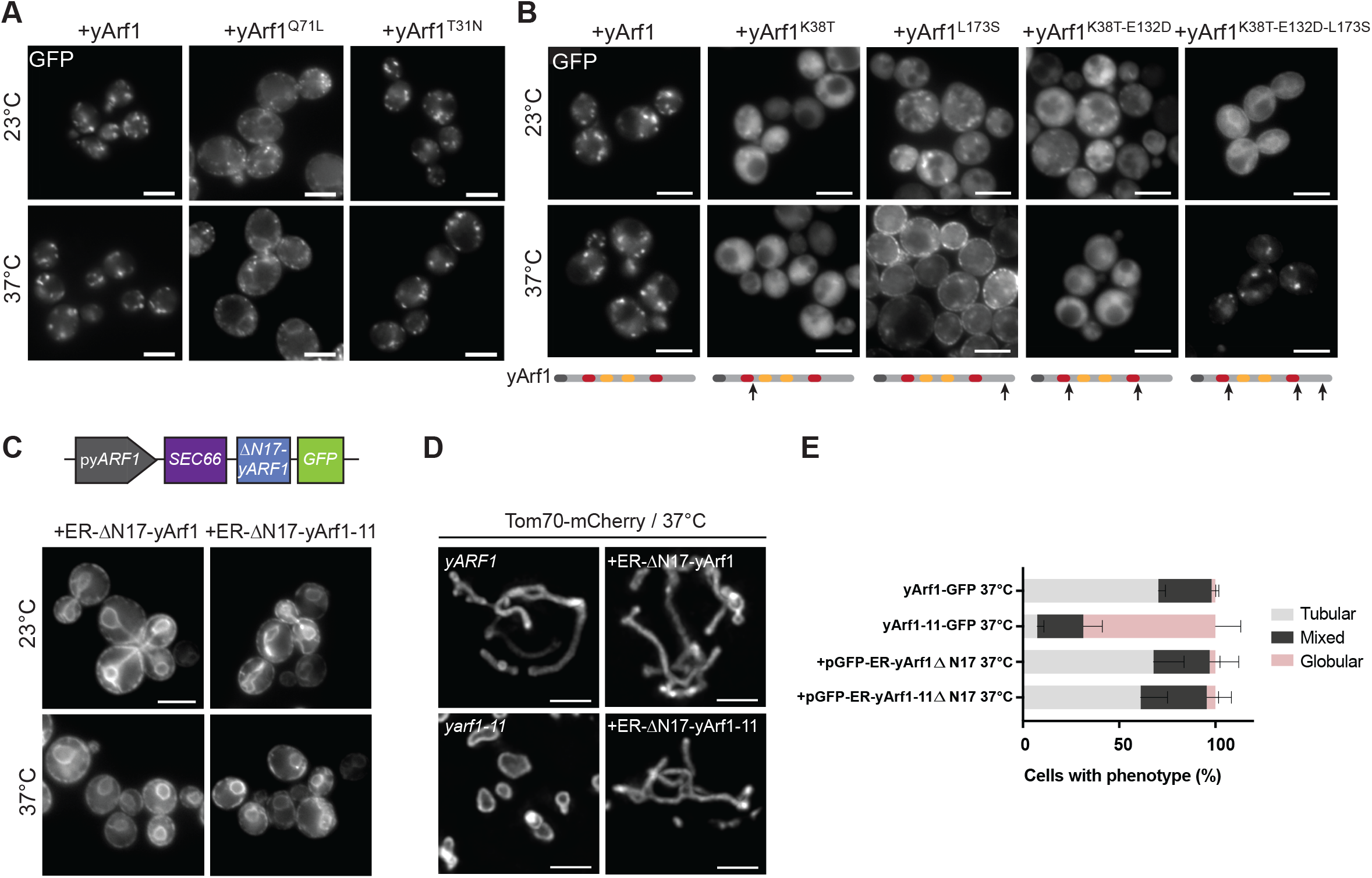
yArf1-11 localization on LD induces mitochondria fragmentation. **A)** Localization of yArf1, constitutively active (Q71L) or constitutively inactive (T31N) forms of yArf1 fused to GFP in *S. cerevisiae* (YPH500) grown at 23°C or shifted to 37°C for 30 min. Constructs were expressed from the centromeric low copy number plasmid pGFP33. **B)** Localization of yArf1, or yArf1 bearing single (K38T, L173S), double (K38T-E132D) or triple (K38T-E132D-L173S) substitutions y*arf1-11* mutations fused to GFP in *S. cerevisiae* (YPH500) grown at 23°C or shifted to 37°C for 30 min. Constructs were expressed from the centromeric low copy number plasmid pGFP33. **C)** Schematic of the construct designed to anchor yArf1 on the ER via Sec66. y*ARF1* deleted in its myristoylation sequence (ΔN17) was expressed from its endogenous promoter and fused to GFP on its 3’ end. Localization of ER-anchored ΔN17-yArf1-GFP or yArf1 bearing yArf1-11-GFP variant in YPH500 cells grown at 23°C and 37°C. **D)** Cells expressing yArf1/11 fused to GFP, or expressing ER-ΔN17-yArf1/11-GFP were grown at 37°C and mitochondria were imaged by fusing Tom70 with Cherry. Scale bar: 2 µm. Mean and standard deviation are shown. **E)** Measurements of mitochondria phenotypes (Tubular, Mixed or Globular) based on images taken in D. Scale bar: 5 µm See also Figure S3

We asked next which mutation, or combination of mutations, is responsible for the localization of yArf1-11 to the ER and LDs by re-introducing yArf1-11 mutations in WT yArf1 fused to GFP (**Table S1, S2**). The single mutations K38T and L173S as well as the K38T-E132D pair perturbed yArf1 localization, but failed to localize yArf1 to the ER at 23°C (**Figure 3B**). Only the reconstitution of all three mutations (K38T-E132D-L173S) caused yArf1 to be on the ER at 23°C and on LDs at 37°C (**Figure 3B**), but none of the combinations tested had a dominant phenotype and impaired WT cell growth (**Figure S3A**). Altogether, this suggests that all three mutations are required to generate a hyperactive form of yArf1 that localizes to the ER at 23°C and to LDs at 37°C.

Since yArf1 is present at mitochondrial fission and fusion sites, but yArf1-11 is sequestered on LDs at 37°C, we wondered whether altered yArf1 localization causes the mitochondrial fragmentation phenotype. To this end, we replaced the N-terminal membrane targeting sequence of Arf1 with the transmembrane domain of the ER protein Sec66 (**Figure 3C**). As expected, at both permissive and restrictive temperatures, the anchored forms yArf1 and yArf1-11 remained on the ER. Their sequestration at the ER did not affect growth even in the absence of any wildtype y*ARF1* (**Figure 3C, Figure S3B, C**). Under these conditions, the mitochondrial network remained tubular in the strain expressing ER-anchored yArf1-11, even when cells were grown at the restrictive temperature (**Figure 3D, E**). Therefore, sequestration of yArf1-11 on lipid droplets results in mitochondrial fragmentation.

### Mammalian Arf1-11 localizes on mitochondria and LD

Since Arf1 also plays a role in mitochondrial dynamics in mammalian cells, we wondered whether the mutation-driven localizations of Arf1 observed in yeast were also conserved in mammalian cells. Therefore, we inserted all three yArf1-11 mutations at their corresponding positions in mammalian Arf1 (mArf1-11), and C-terminally fused WT mammalian Arf1 (mArf1) or the mutant to GFP (**Figure S4A**). These proteins were expressed in HeLa cells in which endogenous m*ARF1* was knocked out by CRISPR/Cas9 (*ARF1* KO; **Figure S4B, C**) [44]. Knock-out of mArf1 only yielded minimal effects on cell proliferation [44]. In contrast, mArf1-11 expression in these cells for more than 3 days led to drastic cell death, while mArf1 expression did not (**Figure S4D**). Thus, like in yeast, the predominantly active form of mArf1-11 has severe effects on cell survival.

As expected, mArf1-GFP was present on the Golgi (co-localization with GM130) and on COPI vesicles (co-localization with βCOP) (**Figure 4A, C, E, Figure S4E**). In contrast, mArf1-11 only modestly localized to the Golgi (**Figure 4B, Figure S4F**), instead mArf1-11 decorated tubular and large round structures. Since yArf1-11 mainly localizes on the ER in yeast at 23°C, we asked whether the tubular fraction of mArf1-11 also colocalizes with the ER marker CLIMP63. However, when compared to mArf1, the mutant co-localized only moderately with CLIMP63 structures (**Figure 4C, D**). It has to be noted that yArf1-11 only localizes to the ER at 23°C and not at 37°C, the growth temperature of mammalian cells. In contrast, the mutant mArf1-11 was found on TOM20-positive tubules where mArf1-positive vesicles were sometimes juxtaposed to mitochondria in agreement with mArf1 function in mitochondria division or transport [26-28], indicating that mArf1-11 localizes to mitochondria in mammalian cells (**Figure 4E, F**). In yeast, yArf1-11 is mainly present on LDs at 37°C. Therefore, we asked whether large round mArf1-11 structures could also be LDs. Staining lipid droplets with the fluorescent fatty-acid BODIPY RedC12 showed that, while mArf1 sometimes juxtaposed with LDs (**Figure 4G**), mArf1-11 was found on the circumference of LDs (**Figure 4H**). Therefore, Arf1-11 LD localization is conserved from yeast to mammals at 37°C.

**Figure 4.**
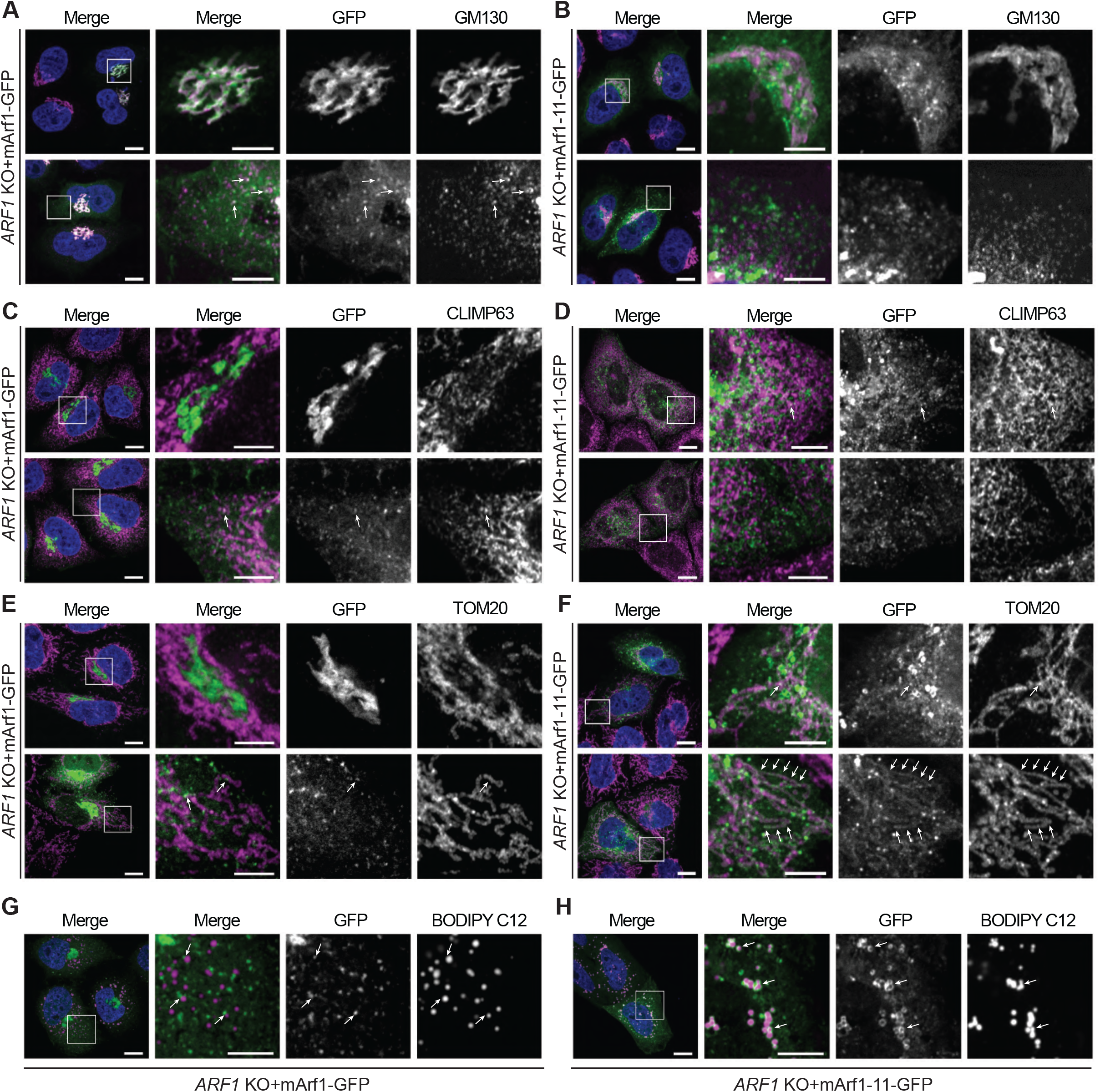
Mammalian Arf1-11 localizes on mitochondria and LD. **A-B)** Mammalian Arf1 (mArf1; **A**) or mArf1-11 (**B**) fused to GFP were expressed in CRISPR/Cas9-mediated *ARF1* knockout HeLa cells (*ARF1* KO). Co-localization with the Golgi was done by immunostaining against the marker GM130. Squares show magnification of a perinuclear and distal portion of the cell. **C-D)** Co-localization of mArf1- (**C**) and mArf1-11-GFP (**D**) expressed in the *ARF1* KO cell line with the ER done by immunostaining against the marker CLIMP63. Squares show magnification of a perinuclear and distal portion of the cell. **E-F)** Co-localization of mArf1- (**E**) and mArf1-11-GFP (**F**) expressed in the *ARF1* KO cell line with mitochondria done by immunostaining against the translocase of mitochondrial outer membrane TOM20. Squares show magnification of a perinuclear and distal portion of the cell. **G-H)** Co-localization of mArf1- (**G**) and mArf1-11-GFP (**H**) expressed in the *ARF1* KO cell line with lipid droplets done by incubation with the fluorescent fatty-acid BODIPY RedC12. Squares show magnification of distal portion of the cell. All images were acquired 24h after transfection. Scale bars: 10 µm and 5 µm (inlays) See also Figure S4

### The hyperactive form of Arf1 induces triacylglycerol accumulation

When we analyzed the yeast *arf1-11* mutant phenotype, we noticed in our TEM pictures that the mutant cells contained more LDs than cells expressing yArf1 both at the permissive and restrictive temperatures (**Figure 2E-H, Figure 5A, Figure S5A**). We confirmed this observation by staining LDs with LipidTox and Nile Red in yeast (**Figure 5B, Figure S5B**) and in mammalian cells (**Figure 5C**), respectively. Lipid droplets are specialized organelles primarily known for their role in energy storage in the form of neutral lipids, mainly triacylglycerol (TAG) and sterol esters (SE). Although lipidomic analysis of the Arf1 and Arf1-11 yeast strains did not reveal any differences in the levels of SE at 23°C or 37°C (**Figure 5D**), we found a strong increase in TAG levels in the y*arf1-11* mutant strain at both temperatures compared to y*ARF1* (**Figure 5E**). Strikingly, these elevated TAG levels were not matched by changes in the overall phospholipid composition (**Figure 5F**). Thus yArf1-11 leads to an increase specifically of TAG in LDs. This TAG accumulation could either be due to altered signalling pathways that would promote LD proliferation or that FA efflux from LDs is perturbed. These possibilities are not mutually exclusive. Nevertheless, reduction of FA efflux from LDs should induce some compensatory mechanisms such as extension of LD contact sites. Indeed, the LDs appeared often to cluster and to be in contact with the ER and/or mitochondria (**Figure 5G**). The expansion in LD number in y*arf1-11* was also concomitant with an increase in ER-mitochondria contacts and contact length in the *arf1-11* strain both at 23°C and 37°C (**Figure 5I, J**). The observation that mitochondria increased their contact-sites with ER and LDs led us to hypothesize that y*arf1-11* might perturb FA flux and as a compensatory mechanism organellar contacts are upregulated.

**Figure 5.**
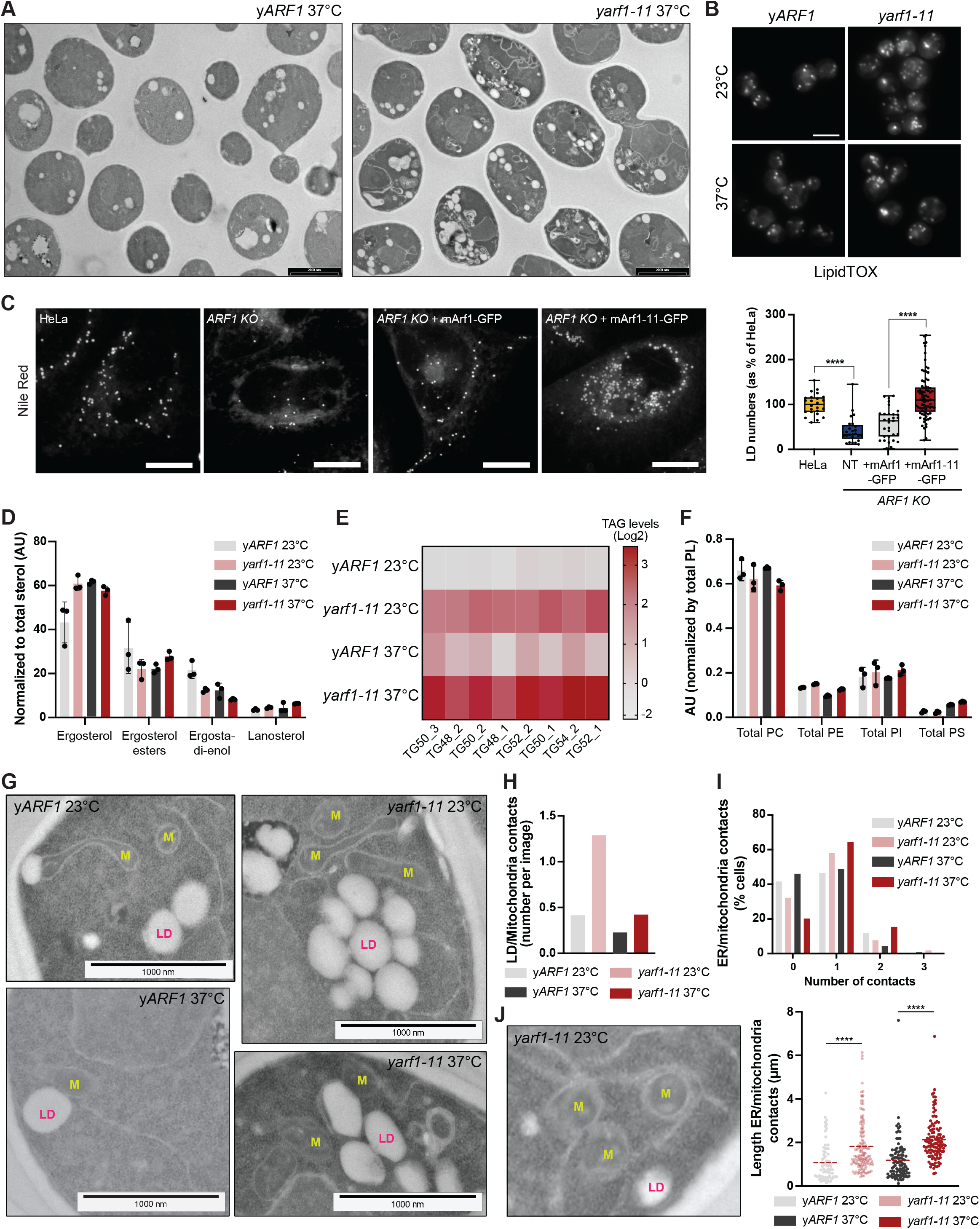
The hyperactive form of Arf1 induces triacylglycerol accumulation. **A)** Transmission electron microscopy of y*ARF1* and y*arf1-11* strains grown at 37°C for 30 min. Scale bars: 2000 nm. **B)** LipidTox staining of LDs in y*ARF1* and y*arf1-11* strains grown at 23°C or 37°C. Scale bar: 5 µm **C-D)** Measurements of sterols and derivatives (**C**), triacylglycerols (TAG; **D**) in the y*ARF1* and y*arf1-11* strains grown at 23°C or 37°C. Levels were normalized to total sterols and are shown as arbitrary units (AU). **E)** Nile Red staining of LDs in parental HeLa cells (Control), *ARF1 KO* HeLa cells, and *ARF1 KO* HeLa cells expressing mArf1 or mArf1-11. For each cell line, the numbers of LD were quantified. Means and standard deviations are shown. Unpaired two-tailed *T*-test. Scale bar: 5 µm. Images were acquired 24h after transfection. **F)** Measurements of phospholipids in y*ARF1* and y*arf1-11* strains grown at 23°C or 37°C. PC: phosphatidylcholine, PE: phosphatidylethanolamine, PI: phosphatidylinositol, PS: phosphatidylserine. Means and standard deviations are shown. **G)** Transmission electron microscopy (TEM) of y*ARF1* and y*arf1-11* strains grown at 37°C for 30 min. LD: lipid droplet; M: mitochondria. Scale bars: 1000 nm. **H)** Quantification of contacts between LDs and mitochondria based on TEM images done in (**G**). **I-J)** Quantification of contacts (**I**) and contact length (**J**) between the ER and mitochondria based on TEM images done in (**G**). The TEM image in (J) shows an example of ER/mitochondria contacts length. Means are shown; each dot represent a single measurement. Unpaired two-tailed *T*-test, \*\*\*\**p* > 0.0001. See also Figure S5.

### yArf1 regulates the first steps of β-oxidation

Therefore, we decided to concentrate on FA metabolism. To be used as energy source, FAs have first to be activated by coenzyme A (CoA) in LDs, which produces acyl-CoA. This activated acyl-CoA can then enter the β-oxidation pathway yielding acetyl-CoA. In contrast to mammalian cells, where FA β-oxidation takes place both in peroxisomes and in mitochondria, yeast β-oxidation is confined to peroxisomes [45, 46]. The produced acetyl-CoA has then to be transported from peroxisomes to mitochondria. However, the precise pathways and mechanisms are still elusive. We assumed that the accumulation of TAG and mitochondrial fragmentation could be due to impaired peroxisome biogenesis, defects in metabolite transport or due to β-oxidation deficiency.

However, we have previously shown that the ER stress response is elevated in y*arf1-11* [23]. Therefore, we first asked whether increase in LD biogenesis was due to ER stress. We deleted in wildtype yeast, the two mammalian *FIT2* homologues, *SCS3* and *YFT2*, known to connect ER stress response and LD biogenesis [47], and to maintain cellular proteostasis and membrane lipid homeostasis at the ER (**Figure S6A, B**) [48]. In these strains, LD accumulation and mitochondrial morphology remained unaffected and did not phenocopy y*arf1-11*. Thus, TAG accumulation and increased LD biogenesis were not a secondary effect due to ER stress in the y*arf1-11* strain.

To test whether peroxisomes were impaired in y*arf1-11* mutants, we first checked peroxisome biogenesis using Pex3 as a marker. We could, however, not detect any defects in the biogenesis of peroxisomes in y*arf1-11* cells (**Figure 6A**). We considered next that the flux of metabolites between peroxisomes and mitochondria might be impaired. Pex34 is part of the tether that organizes peroxisome-mitochondria contact sites [12]. Deletion of *PEX34* in y*ARF1* and y*arf1-11* led to a small increase of cells showing globular mitochondria, with the exception of the y*arf1-11* strain at the restrictive temperature where the phenotype is already at its highest (**Figure 6B**). This suggests impaired metabolite transfer from peroxisomes to mitochondria in y*arf1-11* that might contribute to globular mitochondrial shape. Nevertheless, the change in mitochondrial morphology in *Δpex34* was not as strong as in y*arf1-11*. Either there are additional tethers or peroxisomal acetyl-CoA production might be reduced.

**Figure 6.**
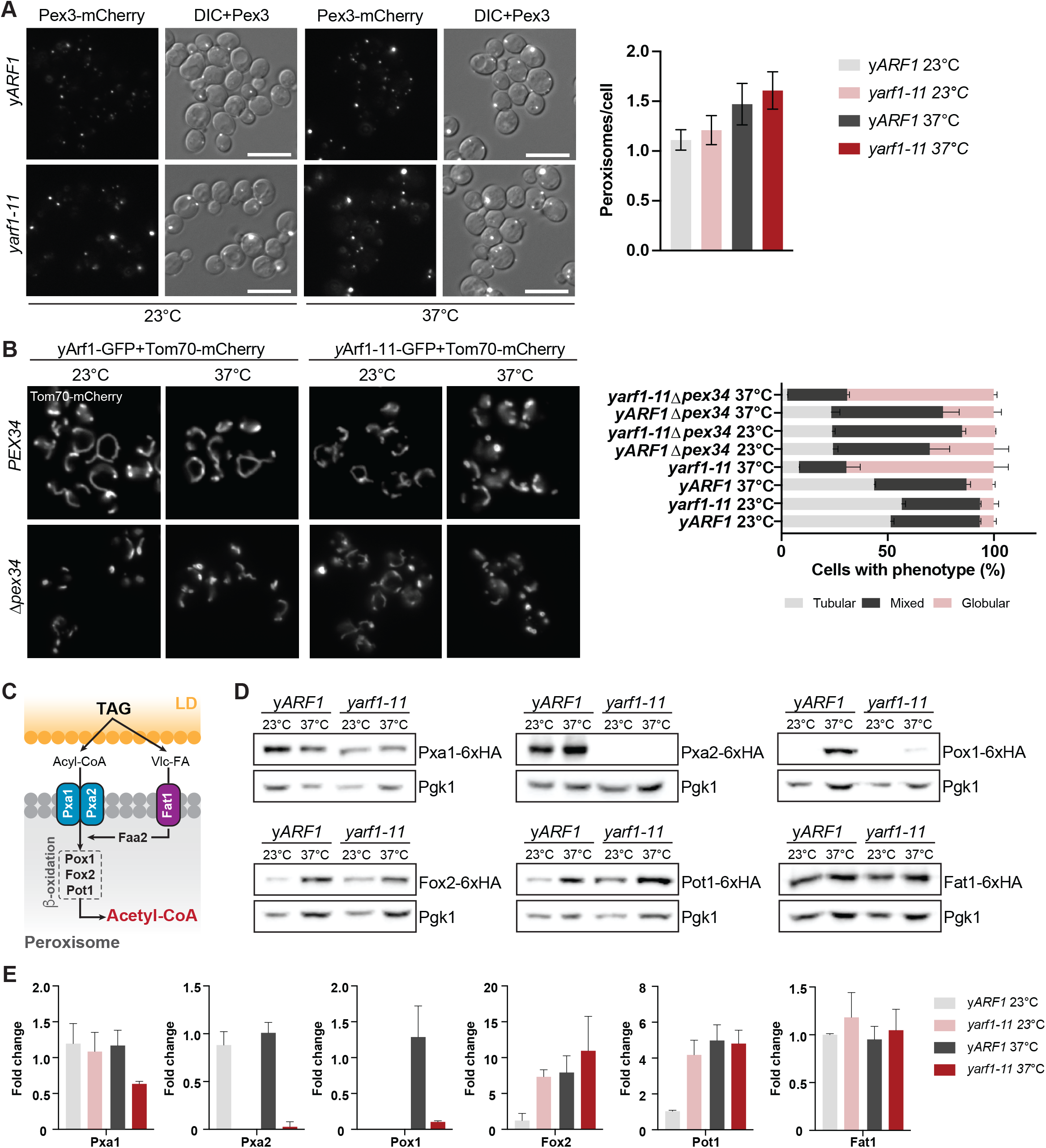
yArf1 regulates the first steps of β-oxidation. **A)** Peroxisome biogenesis followed by microscopy using the peroxisomal marker Pex3 fused to mCherry in the y*ARF1* and y*arf1-11* strains grown at 23°C or 37°C (left panels). Quantification of peroxisomes per cell in each strain and condition (right panel). Means and standard deviations are shown. **B)** Mitochondrial morphology imaged in the y*ARF1* and y*arf1-11* parental strains (*PEX34*) or in strains deprived of *pex34* (Δ*pex34*) grown at 23°C or 37°C. In these strains, Tom70 was fused to mCherry and used as a mitochondrial marker (left panels). Quantification of the mitochondrial phenotypes observed in each strain and condition. Means and standard deviations are shown. **C)** Schematic of triacylglycerol (TAG) mobilization to synthesize acetyl-CoA by peroxisomal β-oxidation in yeast. Relevant proteins monitored in (**D**) are shown. Vlc-FA: very long-chain fatty acids. **D)** Immunoblot analysis of all β-oxidation proteins, both acyl-CoA transporters and the VlcFA transporter genomically fused to 6xHA in the Arf1 and *arf1-11* strains grown at 23°C or 37°C. Pgk1 was used as loading control. **E)** Relative fold changes in protein levels from immunodetections done in (D). Means and standard deviations are shown. Scale bars: 5 µm. See also Figure S6.

Therefore, we determined whether FA β-oxidation was defective in the y*arf1-11* strain (**Figure 6C**). Key components involved in FA β-oxidation were HA-tagged at the genomic level and their protein levels detected in the y*ARF1* and y*arf1-11* strains at 23°C or 37°C by immunoblot. Indeed, the levels of the two subunits of the heterodimeric FA transporter Pxa1 and Pxa2 were affected in y*arf1-11* (**Figure 6D, E**). While we observed an approximately 50% decrease of Pxa1 in the y*arf1-11* strain at the restrictive temperature, Pxa2 was virtually absent in the y*arf1-11* strain, both at 23°C and 37°C. Likewise, the first and rate-limiting enzyme involved in β-oxidation, acyl-CoA oxidase Pox1, was nearly undetectable at 37°C in the y*arf1-11* strain, while its levels were increased in y*ARF1* under the same condition. However, not all peroxisomal proteins were affected in the y*arf1-11* mutant. The level of the very long-chain FA transporter Fat1 and two other enzymes of the β-oxidation cascade, Fox2 and Pot1, were not reduced in y*arf1-11*. Thus, our data indicate that both β-oxidation as well as flux of acetyl-CoA from peroxisomes to mitochondria are impaired in y*arf1-11* at the restrictive temperature.

To corroborate these findings, we tested whether yArf1 is indeed required for FA metabolization. We grew y*ARF1* and y*arf1-11* cells in the presence of saturated FAs (+SFA) or in the presence of cerulenin, where endogenous FA synthesis is abolished (+SFA+Cer) and cells rely on exogeneous FA uptake (**Figure S6C**). In the presence of SFA, y*arf1-11* cells showed growth defects already at 23°C, which was further exacerbated in the presence of cerulenin. The same growth phenotype was observed in y*arf1-11* strains unable to generate TAG (*Δlro1Δdga1*) or with reduced metabolite transfer from peroxisomes to mitochondria (Δ*pex34*), further confirming that FAs metabolization is impaired (**Figure S6C**). Besides FAs, acetate can also be metabolized by yeast cells to produce acetyl-CoA. In the presence of 0.3 M sodium acetate, all strains were able to grow at 23°C, but none of the y*arf1-11* mutant strains sustained growth at 30°C, a semi-permissive temperature (**Figure S6D**). Taken together, our data provide evidence that the y*arf1-11* strain is unable to synthetize acetyl-CoA and that hence yArf1 regulates peroxisome-related metabolism.

### Fatty acid transfer to mitochondria is impaired in y*arf1-11* and leads to mitochondria fragmentation

Our data indicate that in y*arf1-11* cells FAs cannot be efficiently metabolized at the restrictive temperature. Given that metabolites of β-oxidation are used in mitochondria, we suspected that mitochondrial morphology might be affected just by the lack of FAs. To test this hypothesis, we first deleted the two TAG synthases *LRO1* and *DGA1* and determined mitochondrial morphology. As expected, the LD marker Erg6 remained in the ER resulting from a lack of LD biogenesis in all strains and conditions tested, confirming the absence of TAG (**Figure S7A, C**). We noticed an increase in the proportion of cells harbouring globular mitochondria already in the y*ARF1* strain both at 23°C and 37°C, and in the y*arf1-11* at 23°C (**Figure S7B, C**), indicating that TAG removal is sufficient to phenocopy the mitochondrial morphology we observe in the *arf1-11* mutant at the restrictive temperature. To confirm this, we investigated the impact of FA deprivation on mitochondrial morphology by treating cells with cerulenin [49] (**Figure 7A-C**). Treatment of cells with cerulenin for 6 h efficiently reduced the levels of FAs and LDs as seen by Erg6 localization in the ER (**Figure S7D**), and slowed down growth at both temperatures tested (**Figure S7E**). Under these conditions, the proportion of cells with globular mitochondria increased drastically in all strains regardless of the temperature tested (**Figure 7A, B**), confirming that the mitochondrial phenotype we observe in the y*arf1-11* mutant at the restrictive temperature is a consequence of a lack of FA availability. Thus, yArf1 contributes to mitochondrial morphology directly through involvement in mitochondrial dynamics and indirectly by regulating FA metabolism. To corroborate the results above and to show more directly that lipid flux into mitochondria is impaired in y*arf1-11* cells, we followed the transport of the red-fluorescent FA derivative Bodipy C12 (Red-C12) to mitochondria (**Figure 7D**). After 30 min, Red-C12 efficiently reached mitochondria in y*ARF1* cells (**Figure 7E**), while under the same conditions it mainly remained in structures that could be ER and LD and was rarely transferred to mitochondria in y*arf1-11* (**Figure 7E**).

**Figure 7.**
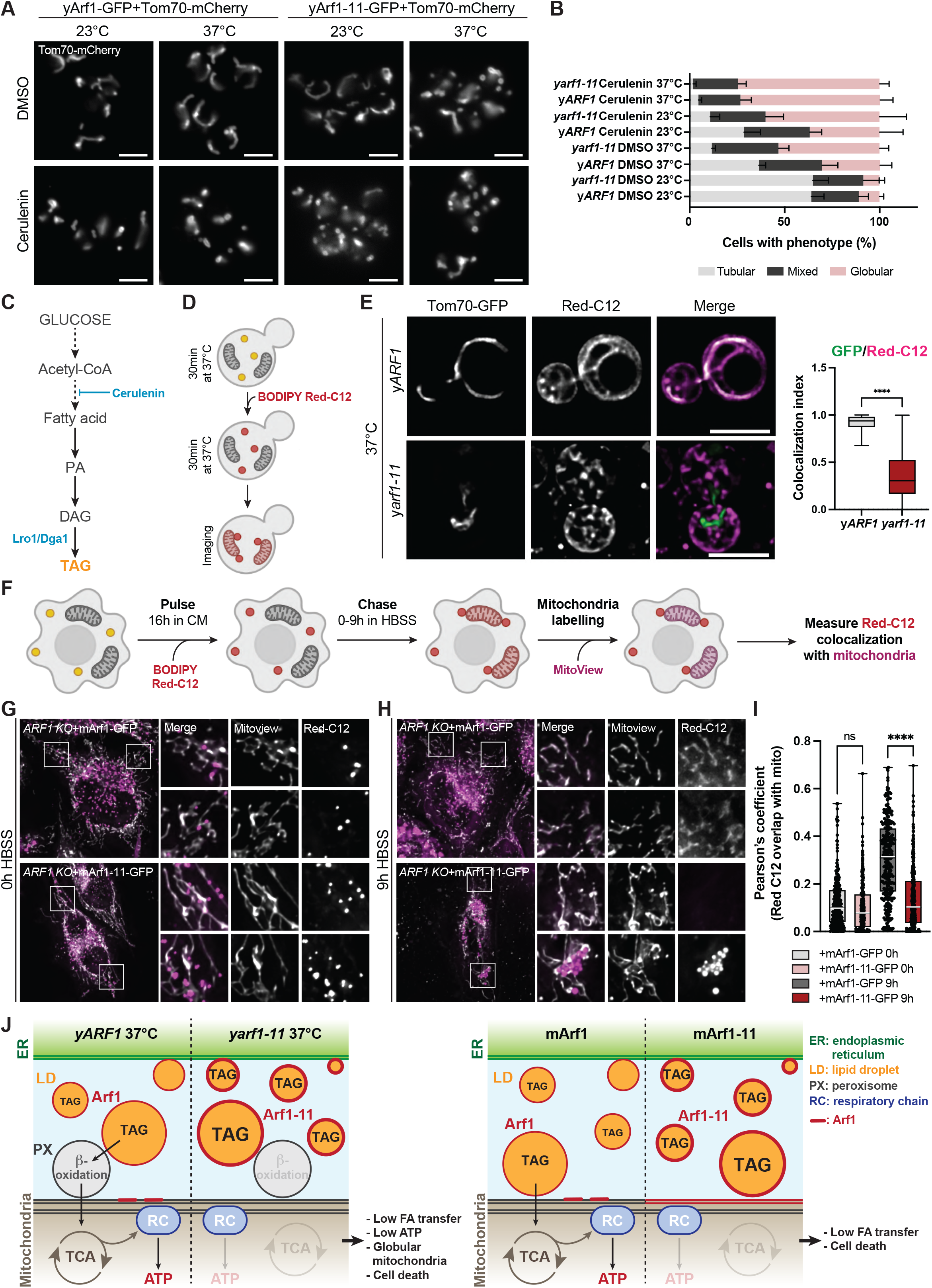
Fatty acids transfer to mitochondria is impaired in y*arf1-11* and lead to mitochondria fragmentation. **A)** Mitochondrial morphology imaged in the y*ARF1* and y*arf1-11* parental strains (*LRO1 DGA1*) or in strains deprived of *LRO1 DGA1* (Δ*lro1* Δ*dga1*) grown at 23°C or 37°C. In these strains, Tom70 was fused to mCherry and used as a mitochondrial marker. **B)** Quantification of the mitochondrial phenotypes observed in (**A**). Means and standard deviations are shown. **C)** Mitochondrial morphology imaged in the y*ARF1* and y*arf1-11* strains grown at 23°C or 37°C and treated with either DMSO or the fatty acid synthesis inhibitor cerulenin. **D)** Quantification of the mitochondrial phenotypes observed in (**C**). Means and standard deviations are shown. **E)** Metabolic pathway leading to triacylglycerol (TAG) synthesis. Fatty acids are used to produce phosphatidic acid (PA), which can be further converted to diacylglycerol (DAG) and to TAG inside LDs by the Lro1 and Dga1 enzymes. **F)** Fatty acid transfer to mitochondria monitored in the yeast *ARF1* and *arf1-11* strains grown at 37°C using the fluorescent fatty acid BODIPY Red-C12. Colocalization of GFP signal (mitochondria) over Red-C12 one (fatty acids) was measured using Mander’s colocalization index. Means and min to max are shown. Unpaired two-tailed *T*-test, \*\*\*\**p* > 0.0001. **G-H)** *ARF1 KO* cells expressing mArf1- or mArf1-11-GFP were pulsed with BODIPY Red-C12 for 16h, incubated 1h in complete medium (CM), transferred in nutrient depleted medium (HBSS) (0h; **G**) and chased HBSS for 9h (**H**). Scale bar: 10 µm. BODIPY Red-C12 was initiated 24h after mArf1 or mArf1-11 transfection. **I)** Relative BODIPY Red-C12 localization measured by Pearson’s colocalization index. Two-way ANOVA using Sidak’s multiple comparison test, \*\*\*\**p* > 0.0001. **J)** Schematic of the model we propose for Arf1 role in FA metabolization and how this affects maintenance of mitochondria morphology. See discussion for more details. Scale bars: 5 µm. See also Figure S7 and S8

### Mitochondrial OXPHOS activity and ATP synthesis are impaired in y*arf1-11*

Mitochondria fragmentation has been described as a general mechanism in response to various types of stress [50], such as ATP synthase inhibition [51], oxidative stress [52], or loss of mitochondrial membrane potential (L4J_m_) [53, 54]. We therefore hypothesized that the lack of metabolite transfer from peroxisomes to mitochondria leads to decreased ATP synthesis and an overall loss of mitochondrial homeostasis.

To test whether mitochondrial respiratory capacity is affected in the y*arf1-11* strain, we grew both y*ARF1* and y*arf1-11* strains in the presence of glycerol, a non-fermentable carbon source that requires a functional respiration to be metabolized. On glycerol-containing plates and in the presence of the ATP synthase inhibitor oligomycin (+Oligo), the y*arf1-11* strain showed impaired growth at 30°C suggesting defects in respiratory chain (RC) function (**Figure S8A**). Taking this into account, we decided to check directly for RC functionality at 23°C and 30°C.

First, we measured the mitochondrial inner membrane potential (L4J_m_) in both y*ARF1* and y*arf1-11* strains. The y*arf1-11* strain exhibited reduced L4J_m_ when grown at 30°C (**Figure S8B**), together with impaired ATP synthesis, but not hydrolysis (**Figure S8B, C**). We confirmed these observations by directly measuring ATP synthesis and hydrolysis rates on purified mitochondria (**Figure S8D, E**), at the cellular level by using a FRET-based ATP nanosensor [55] (**Figure S8F**) and biochemically after 30-, 120- and 360-min incubation at 37°C (**Figure S8G**). In all cases, the outcome was a lower ATP level in y*arf1-11*. The inability of the y*arf1-11* to synthesize ATP was attributed to a decrease in oxygen consumption (*i*.*e* lower respiratory rate; **Figure S8H**) and not due to uncoupled oxidative phosphorylation (P/O; **Figure S8I**). This evidence led us to conclude that the low ATP synthesis rate in y*arf1-11* is a direct consequence of low respiratory rate. Taken together, we revealed a novel role for yArf1 in FA metabolism and as a consequence in respiration and ATP production and mitochondrial morphology.

### Arf1 controls FA flux into mitochondria in mammalian cells

Finally, we tested whether this process was conserved in mammalian cells. We performed a BODIPY Red-C12 pulse-chase assay using *ARF1*-*KO* cells expressing mArf1- or mArf1-11-GFP as reported previously [14]. Cells were pulsed for 16 h in complete media (CM), chased in nutrient-deprived media (HBSS) for 0 or 9 h, and mitochondria were labelled (**Figure 7F**). Under these conditions, FAs were present in LDs both in cells expressing mArf1 or mArf1-11 at the 0 h time point (**Figure 7G**). However, after 9 h of starvation FAs still persisted in LDs and where not transferred to mitochondria in the presence of the mutant mArf1-11, similar to what was observed at the 0 h time point. In contrast, FAs were efficiently transferred in cells expressing mArf1, clearly showing that Arf1 has an evolutionarily conserved role in FA transfer to mitochondria (**Figure 7H, I**). From these experiments we can also conclude that TAG accumulation into LDs in the y*arf1-11* strain and in the *ARF1-KO* cell line expressing mArf1-11 is a direct consequence of impaired FA transfer from LDs to peroxisomes (yeast) and LDs to mitochondria (mammals), which cannot be used for energy production, leading to mitochondrial fragmentation.

## Discussion

We and others have shown previously that Arf1 regulates mitochondrial dynamics and function [26-28]. Here, we elucidate that Arf1 fulfills this role via two independent mechanisms. First, yArf1 is required for both fission and fusion of mitochondria. We provide evidence that yArf1 is present at sites on mitochondria where either fusion or fission occurs. These data reconcile the seemingly inconsistent data from us and others on the mechanism. We and others have shown that Arf1 loss of function resulted in mitochondria hyperfusion in *C. elegans* and in mammalian cells [26, 28]. In contrast, in yeast, the *arf1-11* mutant which yields fragmented mitochondria [26] is a gain-of-function mutant as we determined in this study. Therefore, our data are now all consistent. Second, Arf1 regulates mitochondrial function more indirectly by controlling the flow of FAs and metabolites from LDs to peroxisomes/mitochondria in yeast or to mitochondria in mammalian cells (**Figure 7J**). Therefore, we propose that Arf1 influences mitochondrial morphology and function through at least two independent mechanisms.

A role for Arf1 on peroxisomes and LDs had been established earlier. For peroxisomes it is assumed that Arf1 and COPI are important for peroxisome biogenesis [37-40, 56]. Consistent with this, we did not observe a defect in peroxisome biogenesis with our gain-of-function y*arf1-11* mutant. The peroxisomes appeared, however, to be non-functional in terms of β-oxidation. Our data provide evidence that this deficiency is due to a) strongly reduced FA import into peroxisomes and b) a virtual absence of the acyl-CoA oxidase Pox1. We propose that, because the FA cannot be metabolized in the peroxisomes, they remain stored in LDs. This scenario is supported by the fact that deletion of Pox1 or Pxa1 in yeast leads to increased TAG levels in LDs [57]. It is conceivable, however, that in addition Arf1-11 accumulates in such high concentration on the LDs that together with its effectors such as coatomer, it might produce a shell reducing neutral lipid flow towards peroxisomes (**Figure 7J**). Accordingly, FA efflux from LDs might be decreased. These are not mutually exclusive possibilities. Yet, as a consequence, the transfer of acetyl-CoA to mitochondria is reduced, which leads to depletion of substrate for the TCA cycle and ultimately to a reduction of ATP production, which in turn causes the fragmentation of mitochondria. We propose that the two ways by which Arf1-11 interferes with mitochondrial morphology and function drives cell death. The roles of Arf1 in FA flow from LDs to mitochondria is conserved from yeast to mammals. This is remarkable because the pathways are not identical. While β-oxidation occurs in yeast in peroxisomes, it is also a mitochondrial function in mammalian cells, underscoring the importance of Arf1 in FA metabolism.

In y*arf1-11*, the contacts between mitochondria, lipid droplets, and the ER are increased. It often appeared as if all three organelles came together. One explanation for this finding could be that in y*arf1-11* the number of LDs is increased as well as the contacts to mitochondria. Recently, it was observed that very few, if any, LDs are not connected to the ER [58]. Thus, in principle, the ER might just come along for the ride. We think, however, that this is not very likely, because there is also an increase of the contacts between ER and mitochondria. We favor the possibility that, because FAs cannot be transferred efficiently to peroxisomes, contact site between ER and mitochondria for lipid transfer are expanded, probably as a stress response. Yet, these expanded contact sites might not be functional in terms of FA transfer, since Red-C12 could reach the ER but not mitochondria.

We were able to discover the importance of Arf1 in FA metabolism because of the gain-of-function mutant *arf1-11*. This mutant represents an important tool to understand GTPase function, because it is not dominant active, but is still hyperactive, thereby providing a tool to less drastically affect Arf1 function. Since we observe similar effects of Arf1-11 in yeast and mammalian cells, we assume that mArf1-11 will also be useful in mammalian cells. We hypothesize that corresponding mutations in other Arf/Arl family proteins would have a similar effect.

Strikingly, Arf1 appears to be a Jack of all trades. Historically, Arf1 has been mostly implicated in vesicle pathways within or exiting the Golgi apparatus. We have previously shown that Arf1 is also involved in mRNA transport and metabolism [23]. Moreover, Arf1 is also present on the ER, peroxisomes, LDs, and mitochondria [26-28, 36, 37, 56]. While most small GTPases have mostly rather precise set of effector molecules that they recruit, Arf1 could be much more promiscuous since some of the processes it is involved in probably do not require either COPI or clathrin or its adaptors. It will be interesting to determine the local interactome of Arf1 on the different organelles to be better understand how activated Arf1 changes locally its membrane environment. Certainly, more studies are required to reveal the full repertoire of Arf1 function.

## Material and methods

### Strains, media and plasmids

Yeast strains were either grown in rich media composed of 1% w/v yeast extract, 1% (w/v) peptone, 40 mg/L-adenine, 2% (w/v) glucose (YPD) or 2% glycerol (YPGly), or in synthetic complete medium (HC) composed of 0.17% (w/v) yeast nitrogen base with ammonium sulfate and without amino acids, 2% (w/v) glucose and mixtures of amino acids (MP Biomedicals) depending on the auxotrophies used for selection. Inhibition of fatty acid synthesis was done by treating cells with cerulenin (10 µg/mL; Alexis Biochemicals, Lausen, Switzerland) for 6 h, or with equal volume of DMSO as a control. Solid media contained 2% (w/v) agar. YPD plates with saturated fatty acids (SFA) contained 1% Brij58 (Fluka, Buchs, Switzerland), 0.5 mM palmitic acid and 0.5 mM stearic acid (Sigma). Non-supplemented plates only contained YPD and 1% Brij58. YPD plates containing 0.33 M acetate pH 6 were prepared by adding 3M sodium acetate (Ambion) to 1L sterilized YPD. ATP synthase activity was inhibited by adding 0.5 µg/µL oligomycin to YPGly plates. Cloning into the pGFP33 plasmid were performed using the Gibson assembly kit (NEB). *ARF1* point mutations were generated by Quick-change Mutagenesis kit (NEB) and KLD reactions, primers were generated with the NEBaseChanger website (http://nebasechanger.neb.com).

### Cell culture

*ARF1* KO Helaa cells were established, mycoplasma tested, and described elsewhere [44]. HeLa cells were grown in high-glucose Dulbecco’s modified Eagle’s medium (DMEM, Sigma-Aldrich) with 10% fetal bovine serum (FBS, Biowest), 2 mM L-glutamine, 100 U/mL penicillin G, and 100 ng/mL streptomycin, 1 mM sodium pyruvate at 37°C and 7.5% CO_2_. For transient cell transfections, cells were plated into 6-well plates to reach 70 % confluency the following day and transfected with 1 µg plasmid DNA complexed with Helix-IN transfection reagent (OZ Biosciences).

### Yeast transformation

Three units of OD_600_ of yeast cells were grown in appropriate YPD or HC media to mid-log phase. Cells were spun down and washed in 1 volume of 1x TE and 10 mM LiAc. The pellet was then resuspended in 350 µL of transformation mix (1x TE; 100 mM LiAc; 8.5% (v/v) ssDNA; 70% (v/v) PEG3000), incubated with DNA (PCR product or plasmid) for 1 h at 42°C, spun down (30 sec at 10,000 ×g at RT), resuspended in 100 µl of YPD or HC media and cells were plated onto selective media and incubated at 23°C or 30°C. Genomic tagging was done according to standard procedures [59].

### Hela cell lines survival assay

Cells were seeded in 12-well plates at a density of 5500 cells/well, which was confirmed by re-counting. Every 24h for 6 consecutive days, cells from one well for each cell line were trypsinized, resuspended in PBS complemented with 2% FCS, and GFP fluorescence from 100,000 cells per sample measured by a Fortessa flow cytometer. After 3 days, all cell lines were trypsinized, diluted to 1:10 and transferred into fresh media.

### Microscopy

Fluorescence and DIC images were acquired with an ORCA-flash 4.0 camera (Hamamatsu) mounted on an Axio Imager.M2 fluorescence microscope with a 63x Plan-Apochromat objective (Carl Zeiss, Germany) and a HXP 120 C light source using ZEN 2.6 software. Image processing was performed using OMERO.insight client, and analyzed with the Fiji software. Number and length of contact sites measured on TEM images were done with the Fiji software.

High-resolution images were acquired with an ORCA flash 4.0 cooled sCMOS camera (Hamamatsu) mounted on a FEI-MORE microscope with a 100x U Plan-S-Apochromat objective (Olympus).

In Hela cell lines, mitochondria were stained by immunofluorescence using TOM20 antibody (1:200, Santa Cruz sc-17764) as marker, GM130 antibody (1:1,000, Cell Signalling 12480S) was used as Golgi marker, β-COP (1:500, gift from the Wieland lab) as COPI vesicles marker and CLIMP63 (1:1,000, gift from the Hauri lab) as ER marker. Secondary mouse (1:500) and rabbit (1:500) Alexa-Fluor 568 (Invitrogen) antibodies were used and mounted with Fluoromount-G mounting media (Thermo Fischer) containing DAPI. Images were acquired using a LSM700 Upright confocal laser-scanning microscope with the Zen 2010 software (Zeiss) equipped with a Plan-Apochromat 63×/1.4 oil-immersion objective lens and two photomultiplier tubes.

### Protein extraction and immunoblot analysis

For yeast cells, 10 mL of mid-log grown cultures were lysed at 4°C in breaking buffer containing 50 mM Tris-HCl pH 8, 300 mM NaCl, 0.6% Triton X100, 1 mM DTT, 9 M Urea, supplemented with half-volume of glass beads (0.25-0.5 mm; ROTH). Cell debris and unbroken cells were pelleted by centrifugation 3,000 xg for 5 min at RT. Equal protein concentration were loaded on 12 -15% SDS-PAGE and transferred onto 0.45 µm nitrocellulose membranes (Amersham). Membranes were blocked with TBST (20 mM Tris, 150 mM NaCl, pH 7.6, 0.1% Tween20) with 5% non-fat dry milk for 30 min and incubated with anti-HA primary antibody (1:5,000, Eurogentec 16B12) or anti-Pgk1 primary antibody (1:5,000, Invitrogen clone 22C5D8) over night at 4°C, followed by 2 h-incubation with HRP-conjugated secondary antibody (1:10,000; anti-mouse, Invitrogen 31430) in TBST. Chemiluminescence signals were detected using Immobilon Western HRP Substrate (Millipore) and imaged using a FusionFX (Vilber Lourmat).

Alternatively, prior to Gga2^GAT^ interaction, yeast cells were resuspended in 1 mL of 0.2 M sorbitol, 25 mM KPO_4_ pH 7, 2 mM EDTA, 0.6% Triton X100, 1x Halt proteases inhibitor cocktail (Thermo Scientific), transferred to Corex glass tubes filled with 500 µL glass beads (0.25-0.5 mm; ROTH) and broken 15 min with a vortex at 4°C with 30 sec intervals on ice. Unbroken cells and debris were pelleted at 3,000 xg for 5 min at 4°C and supernatants (SN) were transferred into new 1.5 mL Eppendorf tubes and spun down at 100,000 xg for 30 min at 4°C in a TLA 100-3 rotor. One mL of SN (S100) was saved for each sample and the pellets (P100) were resuspended in 500 µL of Lysis buffer.

Hela cell were lysed in 20 mM Tris-HCl pH7.5, 150 mM NaCl, 10 mM MgCl_2_, 1% Triton X100, 2 mM PMSF, Protease inhibitors, separated by 15% SDS-PAGE and transferred to Immobilon-P PDVF membranes (Millipore). Membranes were blocked with TBST (20 mM Tris, 150 mM NaCl, pH 7.6, 0.1% Tween20) with 3% non-fat dry milk for 1 h and incubated with primary antibody in TBST with 1% milk over night at 4°C: anti-Arf1 (1:2,500, Abnova MAB10011), and anti-actin (1:100,000, Sigma-Aldrich MAB1501). After washing, the membranes were incubated with HRP-conjugated secondary antibody (1:10,000; anti-rabbit, Sigma-Aldrich A0545 or anti-mouse, Sigma-Aldrich A0168) in TBST with 1% milk. Chemiluminescence signals were detected using Immobilon Western HRP Substrate (Millipore) and imaged using a FusionFX (Vilber Lourmat).

### Arf1-Gga2 recruitment

#### Gga2^GAT^ expression and purification

One liter of *E*.*coli* BL21 strains harboring the pGEX4-GST -GGA2^GAT^ were induced with 0.5 mM IPTG and cells were transferred from 37°C to 30°C for 3.5 h. Cells were then pelleted at 4,000 xg at RT for 10 min, resuspended in 20 mL ice-cold 1x PBS/5 mM EDTA buffer (2.7 mM KCl, 1.5 mM KH_2_PO_4_, 137 mM NaCl, 5.6 mM Na_2_HPO_4_, 1.4 mM NaH_2_PO_4_, 5 mM EDTA/NaOH pH 8.0) containing 1 mM PMSF and 1x Halt protease inhibitors. The resuspended cells were lysed by sonication 7 times for 10 sec (50% duty) on ice, cleared at 6,000 xg at 4°C for 30 min. The supernatant was transferred to ultracentrifuge tubes and further cleared at 100,000 x g for 1h at 4°C. To isolate GST-fusion proteins the supernatant was added to 500 µl glutathione Sepharose magnetic beads and incubated for 1h at 4°C under rotation. Glutathione Sepharose beads were spun down at 500 xg for 5 min and washed three times with 15 ml ice-cold 1 x PBS/5 mM EDTA and twice with 1 mL 1 x PBS/5 mM EDTA on a magnetic stand. Bound proteins were eluted by three consecutive treatments with 250 µL of reduced glutathione buffer (20 mM reduced glutathione, 100 mM Tris/HCl pH 8.0) at 4°C for 10 min incubation each time. Supernatants were dialyzed in 2.5 L dialysis buffer (10 mM HEPES/NaOH pH 7.8, 1 mM MgCl_2_, 1 mM DTT and 0.2 mM PMSF) with slow stirring O/N at 4°C. The next day samples were centrifuged for 1 min at 20,000 xg at 4°C to remove precipitates. The supernatants were frozen in liquid nitrogen in aliquots of 80 µg protein and stored at -80 °C.

#### GGA2GAT pre-loading on Glutathione magnetic beads

Purified GST-GGA2^GAT^ was loaded onto glutathione (80 µg/tube). To do so, 200 µL of resuspended glutathione magnetic beads were taken, vortexed for 10 sec and washed twice in 500 and 400 µL of 1x PBS, 0.5 mM EDTA on a magnetic stand. Beads were then resuspended in 200 µL of Lysis buffer (0.2 M sorbitol, 25 mM KPO_4_ pH 7, 2 mM EDTA, 0.6% Triton X100, 1x Halt proteases inhibitor cocktail) and pure GST-GGA2^GAT^ was added. Binding was done for 30 min on a rotating stand at 4°C, followed by one wash in 300 µL of Lysis buffer, and final resuspension in 400 µL of Lysis buffer.

#### Binding and elution

S100 and P100 fractions of yeast cells were incubated with pre-bound GGA2^GAT^ on beads for 1h on a rotating wheel at 4°C. Washes were done 3x in 50 µL Lysis buffer and elution was done by adding 30 µL of Laemmli 2x to the beads and 5 min incubation at 95°C. For each gel, 10 µL/lane was used for the lysis, FT and elution samples.

### Transmission Electron Microscopy (TEM)

Cells were grown to mid-log phase and fixed into the YPD media with 0.2% Glutaraldehyde and 3% Formaldehyde final concentration O/N at 4°C. The next day, cells were pelleted and washed 3 times with 0.1 M HEPES buffer pH 7 and incubated for 30 min in 1% NaJO_4_ in HEPES buffer, washed 3 times, and free aldehydes were quenched with 50 mM NH_4_Cl in HEPES buffer for 30 min. After that, pellets were dehydrated through a series of methanol (50%, 70%, 90%) then infiltrated with LR Gold resin (Polysciences) according to the manufacturer’s instructions and allowed to polymerize at -10°C under an UV lamp for 24 h. Sections of 60-70 nm were collected on carbon-coated Formvar-Ni grids. Sections were blocked with PBST (PBS+ 0.05% Tween 20) + 2% BSA for 15 min and incubated 3h at RT with anti-GFP antibody (1:100, 6556 Abcam) in PBST/BSA. Sections were washed 5x 5 min with PBS and incubated with goat anti-rabbit secondary antibody (BBI) coupled to 10 nm gold 1:100 in PBST/BSA for 2 h, washed 5x 5 min with PBS and 3x 2 min H_2_O, and stained for 10 min in 2% Ur acetate and 1 min in Pb citrate (Reynold’s solution). Sections were viewed with a Philips CM100 electron microscope.

### Lipid extraction from yeast cell pellets

Lipid extraction and analysis were performed in triplicate. Frozen cell pellets (5×10^8^ cells) were taken from -80°C and 500 µL glass beads were added, then 20 µL of yeast internal standard mix and 20 µl cholesterol were added (see below), followed by 600 µL H_2_O, 1.5 ml methanol, then vortexed for 1 min. 0.75 mL chloroform was added and the samples were vortexed at high speed for 6 min. The solution was transferred to a 13×100mm screw capped glass tube and the beads were washed with 0.6 mL chloroform:methanol (1:2). Following vortexing, the solution was combined with the first extraction. 0.4 mL H_2_O was added and mixed by vortexing. Samples were centrifuged for 10 min at 4,000 rpm (3220 xg) and most of the aqueous phase was removed, followed by transfer of the organic phase to fresh 13×100 tubes without taking interface. 0.4 mL of artificial upper phase (Chloroform/Methanol/Water (3:48:47, v/v/v)) was added and mixed by vortexing. After centrifugation for 10 min at 4000 rpm and removal of most of aqueous phase, the organic phase was transferred to ms vials. 0.3-0.4 mL were transferred to a vial insert (filled up) for analysis using the Thermo Q-Exactive Plus. The extracts in inserts were dried in the Centrivap (Labconco) and the extracts in vials under flow of N_2_.

| Internal Standard | nmole/sample |
| --- | --- |
| PC31:1 | 7.5 |
| PE31:1 | 7.5 |
| PI31:1 | 6 |
| PS31:1 | 4 |
| CL56:0 | 4 |
| C17Cer | 1.2 |
| C8GC | 2 |
| Cholesterol | 20 |

### Sterol analysis by GC-MS

Lipid extracts from approximately 40% of the total sample was dissolved in 0.3 mL Chloroform:Methanol (1:1) and 5 µL was loaded onto a GC-MS (Varian 320 MS) fitted with a fused silica capillary column (15 m*0.32 mm ID, Macherey Nagel ref. 726206.15). The GC program started at 45°C for 4 min, then a gradient to 195°C at 20°C/min, to 230°C at 4°C/min, to 320°C at 10°C/min, to 350°C at 6°C/min followed by a return to 45°C at 100°C/min. Data was collected in the centroid mode. The different sterol species were identified by their retention times and mass spectra using the NIST database and ergosterol and cholesterol standards. Peaks were integrated and amounts calculated with correction for the yield of the internal standard, cholesterol, and using standard curves of cholesterol and ergosterol concentrations.

### Mild base treatment to enrich for sphingolipids

Lipid extracts from approximately 20% of the sample were dried and treated with methylamine and desalted with butanol as described previously [60]. Samples were dried under a flow of N_2_.

### Lipid analysis by electrospray MS using the TSQ-Vantage

Lipid analysis using nanoflow infusion (Advion Nanomate) and multiple reaction monitoring (TSQ Vantage, Thermo) was performed as described [60]. All runs were performed in duplicate with all transitions measured 3 times for a total of at least 6 measurements for each lipid species.

### TAG analysis using the UHPLC-MS using the Q-Exactive plus

Dried samples were resuspended by sonicating in 100 µL of LC-MS-grade chloroform:methanol (1:1, vol/vol). Reversed-phase UHPLC-HRMS analyses were performed using a Q Exactive Plus Hybrid Quadrupole-Orbitrap mass spectrometer coupled to an UltiMate 3000UHPLC system (Thermo Fisher Scientific) equipped with an Accucore C30 column (150 × 2.1 mm, 2.6 µm) and its 20mm guard (Thermo Fisher Scientific). Samples were kept at 8°C in the autosampler, 10 µL were injected and eluted with a gradient starting at 10% B for 1 min, 10-70% B in 4 min, 70-100% B in 10 min, washed in 100 % B for 5 min and column equilibration for an additional 3 min. Eluents were made of 5 mM ammonium acetate and 0.1% formic acid in water (*solvent A*) or in isopropanol/acetonitrile (2:1, v/v) (*solvent B*). Flow rate and column oven temperature were respectively at 350µl/min and 40°C. The mass spectrometer was operated using a heated electrospray-ionization (HESI) source in positive polarity with the following settings: electrospray voltage: 3.9 KV (+); sheath gas: 51; auxiliary gas: 13; sweep gas: 3; vaporizer temperature: 431°C; ion transfer capillary temperature: 320 °C; S-lens: 50; resolution: 140,000; m/z range: 200-1000; automatic gain control: 1e6; maximum injection time: 50 ms. The following setting was used in HCD fragmentation: automatic gain control: 2.5e5; maximum injection time: 120 ms; resolution: 35,000; (N)CE: 30. Xcaliburv.4.2 (Thermo Fisher Scientific) was used for data acquisition and processing. Major TAG species were quantified after removing background and normalization, and was presented as ratio to sample no. 1, respectively.

### Mitochondria fusion and fission dynamics

Yeast cells were grown to mid-log phase in YPD media, and switched to 37°C for 30 min when indicated. Movies were acquired either with an Axio Imager.M2 (Arf1 23°C-37°C and Arf1-11 at 37°C) or the FEI-MORE (Arf1-11 at 23°C), over a period of 2 minutes with sequential image acquisition. Images from the FEI-MORE were further deconvolved in standard mode using the Huygens Pro software. Movies were assembled with the Fiji software and single images prepared on the OMERO.insight client.

### Fluorescent fatty acid pulse-chase

Yeast cells were grown to mid-log phase in HC complete media at 23°C, then switched to 37°C for 30 min. Following this, 1.5 ×10^6^ cells were incubated with 50 µM BODIPY 558/568 C_12_ (Life Technologies) for 30 min at 37°C, washed twice in 1x PBS, 7.5 µM BSA, and resuspended in 20 µL 1x PBS, 7.5 µM BSA supplemented with 20 µL Trypan blue. Images were taken with the FEI-MORE microscope, and deconvolved in standard mode using the Huygens Pro software.

FA pulse-chase experiments in HeLa cells were performed according to Rambold et al. [14]. In short, transfected cells were plated into 8-well imaging chambers (Miltenyi) and incubated for at least 6 hours to let the cells adhere. Then their complete growth medium (CM) was replaced with CM containing 1 µM BODIPY™ 558/568 C12 (Invitrogen) and incubated overnight. Cells were washed three times with CM, incubated for 1 hour in CM and then chased for 9 hours in CM or in HBSS. Mitochondria was labeled with 200nM MitoView Fix 640 (Biotinum) for 20 minutes prior to imaging. Just prior to imaging, CM or HBSS was replaced with imaging buffer containing 25 mM Dextrose supplemented with 10 % FBS or 5 mM Dextrose, supplemented with 0.2 % FBS respectively.

Cells were imaged at 37°C using an inverted Axio Observer microscope (Zeiss) with a Plan Apochromat N 63×/1.40 oil DIC M27 objective and a Photometrics Prime 95B camera. Filters with standard specifications for GFP, dsRed and Cy5 were used to image EGFP, Bodipy 558/568 and Mitoview 640 respectively.

### Lipid droplets staining

Neutral lipids of mid-log grown cells in YPD media were stained with 1µL of the lipophilic fluorophore ReadiStain Lipid Green (InVivo Biosystems) for 30 min at 23°C and 37°C. Cells were then washed 3 times in HC complete media and prepared for microscopy.

Transfected HeLa cells were plated into 8-well imaging chambers (Ibidi, ibiTreat µ-Slide) the day before imaging to reach 50–70% confluency the following day. Just prior to imaging, cells were rinsed with pre-warmed PBS and replaced with imaging buffer (4.5 g/l Dextrose, 1 mM CaCl_2_, 2.7 mM KCl, 0.5 mM MgCl_2_ in PBS supplemented with 0.2 % FBS) containing 400 ng/ml NileRed (Sigma) and incubated for 10 min before starting imaging. The dye was present during imaging. Confocal images were acquired at 37°C with Olympus Fluoview FV3000 system, using an UPLSAPO 60x/1.30 objective with silicone oil, resulting in a xy pixel size of 0.1 µm. Laser intensities were at 0.5–3% for both 488 (GFP) and 561 (DsRed) wavelengths. Sampling speed was 8.0 µs/pixel with a zoom factor of 2.0. All images for corresponding experiments were processed with the same settings to insure comparable results.

### ATP measurements

#### FRET measurement

ATP levels were measured in single cells grown to mid-log phase using a FRET based nanosensor expressed from a cen/ars plasmid, pDR-GW AT1.03YEMK, (Addgene # 28004) similarly to what has been described in [61]. Images were acquired with the Axio Imager.M2, using dedicated CFP, YFP and FRET CFP-YFP filters. ROIs were measured with the Fiji software.

#### Biochemical assay

ATP quantitation was performed using the BacTiter-Glo Cell viability assay (Promega) following manufacturer’s instructions with slight modifications. In brief, cells were resuspended in TE buffer pH7 containing 0.7M sorbitol and 10^6^ cells were used for each assay. Luminescence was measured right after mixing cells with BacTiter-Glo buffer in 96x flat-white plates (Greiner) on a Tecan Infinite M1000Pro with 10 sec orbital shaking (2mm wide, 350 rpm).

### Mitochondrial activity measurements

Mitochondria were isolated from wild type and mutant strains grown for 5-6 generations in 2 L of YPGly at 23 or 28°C by enzymatic method of [62]. The cultures contained 2-5% of ρ^−^/ρ° cells. The values reported are averages of two biological repetitions. Respiratory, ATP synthesis activities and the variations of inner membrane potential were measured using freshly isolated, osmotically protected mitochondria buffered at pH 6.8. The oxygen consumption was measured in an oxygraph with Clarck electrode (Heito, France) at 28°C in thermo-stabilized cuvette. Reaction mixes for assays contained 0.15 mg/ml of mitochondria, 4 mM NADH, 150 µM ADP, 12.5 mM ascorbate (Asc), 1.4 mM N,N,N,N,-tetramethyl-p-phenylenediamine (TMPD), 4 µM CCCP. The rates of ATP synthesis were determined under the same experimental conditions in the presence of 750 µM ADP; aliquots were withdrawn from the oxygraph cuvette every 15 seconds and the reaction was stopped with 3.5% (w/v) perchloric acid, 12.5 mM EDTA. The ATP in samples was quantified using the Kinase-Glo Max Luminescence Kinase Assay (Promega) and a Beckman Coulter’s Paradigm Plate Reader. Variations in transmembrane potential (ΔΨ) were evaluated in the respiration buffer containing 0.150 mg/mL of mitochondria and the Rhodamine 123 (0.5 µg/mL), with A_exc_ of 485 nm and A_em_ of 533 nm under constant stirring using a Cary Eclipse Fluorescence Spectrophotometer (Agilent Technologies, Santa Clara, CA, USA) [63]. For the ATPase assays, mitochondria kept at –80°C were thawed and the reaction performed in absence of osmotic protection and at pH 8.4 according to [64].

### Statistical analysis

All experiments were performed at least in three independent replicates. Unpaired two-tailed *T*-test were calculated for each experiment. When not stated otherwise, means and standard deviations (SD) are shown.

## Acknowledgements

We thank A. Gil and H. Wechlin, S. Begum and S. Feng for excellent technical assistance with some of the experiments. This work was supported by grants of the Swiss National Science Foundation (310030B_163480 and 310030_185127) to AS and the University of Basel, Swiss National Science Foundation, NCCR Chemical Biology and the Leducq Foundation to HR, Swiss National Science Foundation (31003A-182519) to MS, and the National Science Center of Poland (2018-31-B-NZ3-01117) to RK.

## Data availability statement

All data generated or analysed during this study are included in this published article (and its supplementary information files).

## Authors contributions

LE and AS conceived the project. LE performed all yeast cell biology experiments. MP and VS performed the experiments in mammalian cells. IR and HR did the lipidomics analyses. AW and RK carried out the biochemical analyses on yeast mitochondria. The EM experiments were executed by CPB. Data were analyzed by LE, RK, HR, MS and AS. LE and AS wrote the manuscript with input from all authors.

## Movie legends (related to Figure 1)

**Movie 1:** Mitochondria fusion and fission monitored in yArf1-GFP expressing strain together with the mitochondrial protein Tom70 fused to mCherry at 23°C. White arrows indicate sites of fission and yellow arrows fusion.

**Movie 2:** Mitochondria fusion and fission monitored in yArf1-GFP expressing strain together with the mitochondrial protein Tom70 fused to mCherry at 37°C. White arrows indicate sites of fission and yellow arrows fusion.

**Movie 3:** Mitochondria fusion and fission monitored in yArf1-11-GFP expressing strain together with the mitochondrial protein Tom70 fused to mCherry at 23°C. White arrows indicate sites of fission and yellow arrows fusion.

**Movie 4:** Mitochondria fusion and fission monitored in yArf1-11-GFP expressing strain together with the mitochondrial protein Tom70 fused to mCherry at 37°C. White arrows indicate sites of fission and yellow arrows fusion.

## Supplementary figure legends

**Supplementary Figure 1 (related to Figure 1 and 2). yArf1-11 is a hyperactive mutant that localizes to the ER and lipid droplets**

**A)** y*ARF1* and y*arf1-11* strains phenotypes followed by microscopy after 0-, 30-, 60- and 120 minutes incubation time at 37°C.

**B-C)** Co-localization of yArf1-GFP **(B)** and yArf1-11-GFP **(C)** with the *cis*-Golgi marker Mnn9-mCherry grown at 23°C and 37°C. Arrows indicate sites of co-localization between the yArf1 and Mnn9.

**D-E)** Co-localization of yArf1-GFP (**D**) and yArf1-11-GFP (**E**) with the *trans*-Golgi marker Sec7-mCherry and Tvp23-mCherry grown at 23°C and 37°C. Arrows indicate sites of co-localization between the yArf1 and the corresponding markers.

**F-G)** Transmission electron microscopy of the yArf1-11-GFP strain grown either at 23°C (**F**) or shifted at 37°C for 30 min (**G**). yArf1-11-GFP localizations were highlighted by immunogold labelling, and dotted squares shows enlargements of specific localizations.

**H)** Purification steps of the Gga2^GAT^ domain fused to GST. CE: crude bacterial extract; 12k: extract after 12,000 *xg* centrifugation; 100k: extract after 100,000 *xg* centrifugation; FT: flow-through extract unbound on glutathione Sepharose beads; E1-3: three consecutive elutions with reduced glutathione; Pure: Pooled E1-3 fractions after dialysis.

Scale bars: 5 µm.

Scale bars: 500 nm (TEM).

**Supplementary Figure 2 (related to Figure 1). yArf1 regulates mitochondria fusion and fission**

**A-D)** Single time-point images of movies done with strains expressing yArf1-GFP (**A**) or yArf1-11-GFP (**B**) and the mitochondrial protein Tom70-mCherry at 23°C, or at 37°C for yArf1-GFP (**C**) and yArf1-11-GFP (**D**). Merge, together with individual GFP and mCherry channels are shown.

Scale bars: 5 µm

**Supplementary Figure 3 (related to Figure 3). yArf1-11 localization on LD induces mitochondria fragmentation**

**A)** Growth assay of the YPH500 strain bearing the empty pGFP3 vector (+EV), single (K38T, L173S), double (K38T-E132D) or triple (K38T-E132D-L173S) y*arf1-11* mutations fused to GFP on rich YPD plates incubated at 23°C, 30°C or 37°C.

**B)** Growth assay of the ER-anchored ΔN17-yArf1-GFP or Arf1 strains bearing y*arf1-11* mutations (Arf1^K38T-E132D-L173S^) on rich YPD plates or synthetic media lacking uracil (HC -Ura) incubated at 23°C, 30°C or 37°C.

**C)** Growth assay of the ER-anchored ΔN17-yArf1-GFP or yArf1 bearing y*arf1-11* mutations (Arf1^K38T-E132D-L173S^) in YPH500 cells lacking y*ARF1* (Δ*arf1*) on rich YPD plates or synthetic media lacking uracil (HC -Ura) incubated at 23°C, 30°C or 37°C.

**Supplementary Figure 4 (related to Figure 4). Mammalian Arf1-11 localizes on mitochondria and LD**

**A)** Alignment of mammalian Arf1 (mArf1), yeast Arf1 (yArf1) and the yeast mutant *arf1-11* (yArf1-11) amino acids sequences. The Arf1-11 mutations and their corresponding amino acids in mArf1 are showed.

**B)** Immunoblot analysis of Arf1 presence in parental HeLa cells and the CRISPR-Cas9 ARF1 KO cells. For each cell line, three independent biological replicates were analyzed on the same gel.

**C)** Immunoblot analysis of Arf1-GFP presence in *ARF1* KO HeLa cells transfected either with an empty vector (EV), with mArf1-GFP or with mArf-11-GFP. For each cell line, three independent biological replicates were analyzed on the same gel. Actin was used as internal control.

**D)** Cell viability assay represented as percent of GFP-positive cells in the total population after transfection of EV, mArf1-GFP or mArf-11-GFP. Means and standard deviations are shown.

**E-F)** Co-localization of mArf1- (**E**) and mArf1-11-GFP (**F**) expressed in the *ARF1* KO cell line with COPI vesicles done by immunostaining against the coatomer subunit beta (bCOP). Squares show magnification of a perinuclear and distal portion of the cell. Scale bars: 10 µm and 5 µm (inlays). Images were acquired 24h after transfection.

**Supplementary Figure 5 (related to Figure 5). The predominantly-active form of yArf1 induces triacylglycerol accumulation**

**A)** Transmission electron microscopy of y*ARF1* and y*arf1-11* strains grown at 23°C. Scale bars: 2000 nm.

**B)** Quantification of LDs per cell from Figure 5B after LipidTox staining in y*ARF1* and y*arf1-11* grown at 23°C and 37°C.

**Supplementary Figure 6 (related to Figure 6). yArf1 regulates the first steps of β-oxidation**

**A)** Lipid droplets (Erg6) and mitochondria (Tom70) morphologies imaged in the y*ARF1* and y*arf1-11* parental strains (*-*), in strains deprived of *PET10* (Δ*pert10*), *LDH1* (Δ*ldh1*) or *SCS3 YFT2* (Δ*scs3* Δ*yft2*) grown at 23°C or 37°C.

**B)** Schematic of the localizations and roles of the gene candidates studied in (**A**).

**C)** Growth test of parental y*ARF1* and y*arf1-11* strains, deprived of the *LRO1/DGA1* (Δ*lro1* Δ*dga1*) or of *PEX34* (Δ*pex34*), on rich media (YPD), rich media containing lacking both saturated fatty acids (SFA) and cerulenin (Cer) (non-sup), containing SFA or both SFA and cerulenin at 23°C.

**D)** Growth test of parental y*ARF1* and y*arf1-11* strains, deprived of the *LRO1/DGA1* (Δ*lro1* Δ*dga1*) or of *PEX34* (Δ*pex34*), on rich media (YPD) or media supplemented with 0.3M sodium acetate as an acetate source at 23°C and 30°C.

**Supplementary Figure 7 (related to Figure 7). Fatty acids transfer to mitochondria is impaired in y*arf1-11* and lead to mitochondria fragmentation**

**A)** Lipid droplets morphologies and Erg6 localization imaged in the y*ARF1* and y*arf1-11* parental strains (*LRO1 DGA1*) or in strains deprived of *LRO1 DGA1* (Δ*lro1*Δ*dga1*) grown at 23°C or 37°C.

**B)** Mitochondrial morphologies imaged in the parental or Δ*lro1*Δ*dga1* y*ARF1* and y*arf1-11* strains grown at 23°C or 37°C.

**C)** Quantification of the mitochondrial phenotypes observed in (**B**). Means and standard deviations are shown.

**D)** Lipid droplets morphologies and Erg6 localization imaged in the y*ARF1* and y*arf1-11* strains grown at 23°C or 37°C and treated with either DMSO or the fatty acid synthesis inhibitor cerulenin for 6h.

**E)** Quantification of cell population in **D**. Scale bars: 5 µm.

**Supplementary Figure 8. Mitochondrial OXPHOS activity and ATP synthesis are impaired in y*arf1-11***

**A)** Growth test of y*ARF1* and y*arf1-11* strain on rich media containing 2% glucose (YPD), on respiratory media containing 2% glycerol (YPGly) or on respiratory media containing 2% glycerol supplemented with the ATP synthase inhibitor oligomycine (YPGly+Oligo) done at 23°C and 30°C.

**B-C)** Membrane potential measured on isolated mitochondria by Rhodamine 123 at 23°C or 30°C in y*ARF1* and y*arf1-11* strains in the presence of external ADP (**B**) or to test ATP-driven proton pumping in the presence of ATP (**C**).

**D-E)** ATP synthesis (**D**) and ATPase rate (**E**) measured on isolated mitochondria from y*ARF1* and y*arf1-11* strains grown at 23°C or 30°C. Unpaired two-tailed *T*-test, \*\*\**p*= 0.0001, \**p*= 0.0169

**F)** Detection and quantification of intracellular levels of ATP in y*ARF1* and y*arf1-11* strains grown at 23°C or 37°C using a FRET-based ATP-dependent nanosensor. CFP/YFP FRET ratio were measured and used as relative ATP levels. Medians and individual values are showed. Unpaired two-tailed *T*-test, \*\*\*\**p* > 0.0001. n=419-442 cells from 3 independent experiments.

**G)** Relative ATP levels (RLU) measured by Luciferase assay in y*ARF1* and y*arf1-11* strains incubated for 30 min, 120 min and 360 min at 37°C. Means and standard deviations are shown.

**H)** Respiration rate measured on isolated mitochondria from y*ARF1* and y*arf1-11* strains grown at 23°C or 30°C. Two-way ANOVA using Sidak’s multiple comparison test. \*\*\*\**p* < 0.0001, \**p* = 0.0186.

**I)** ATP synthesis coupling to oxygen respiration (P/O) was measure on isolated mitochondria. Percentage of Rho 0 or Rho minus cells were measured.

**Table 1.**
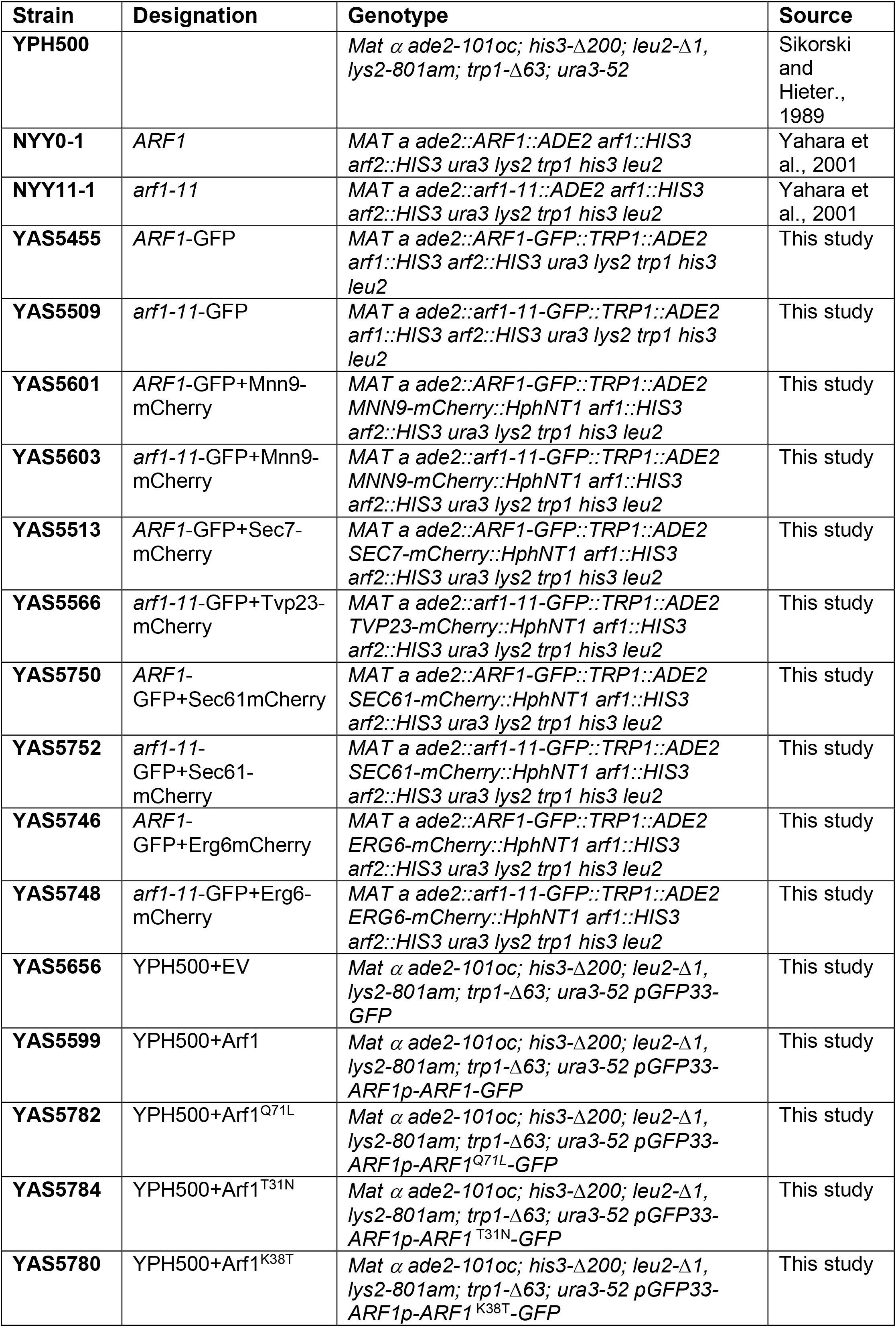
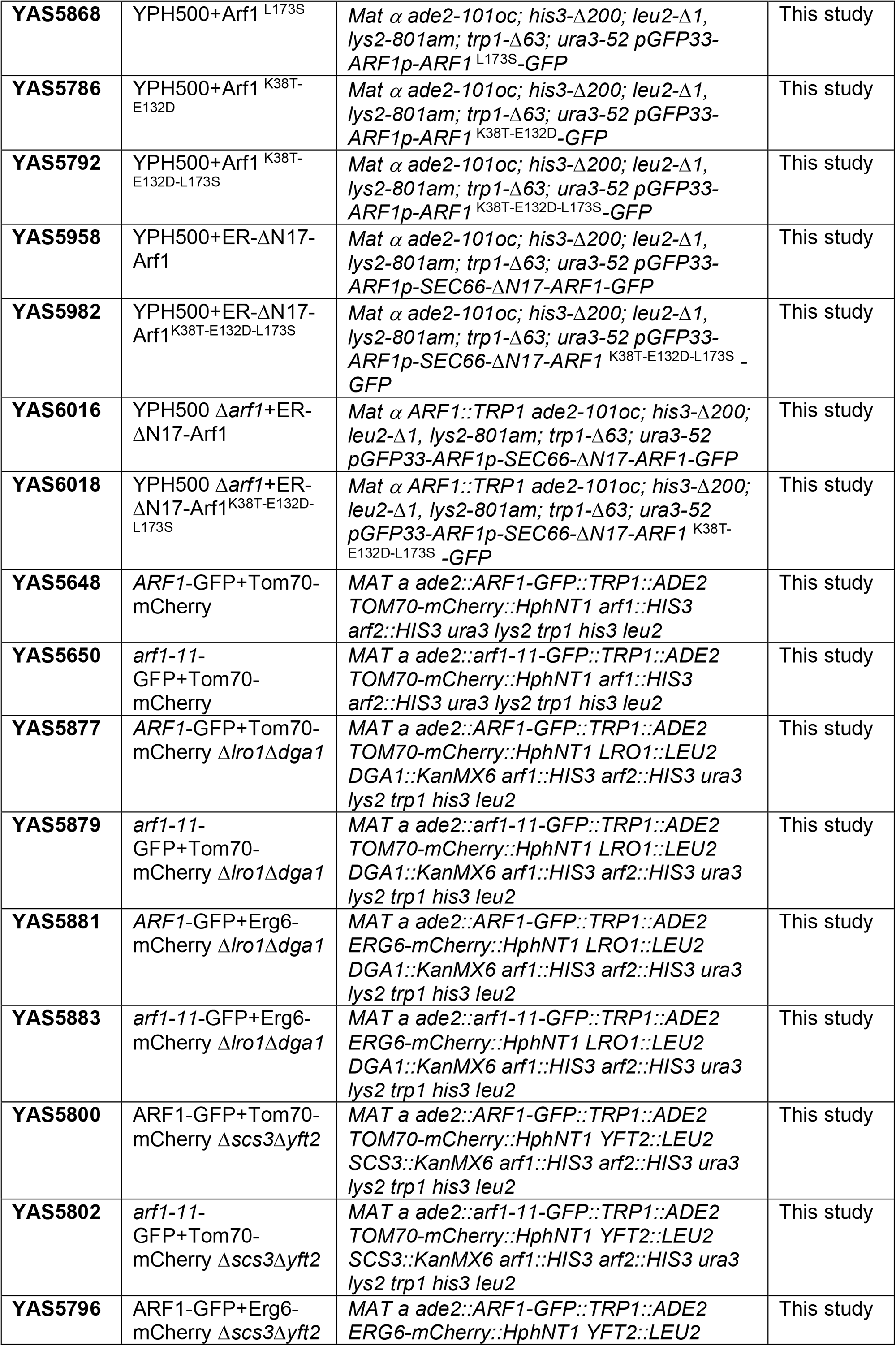
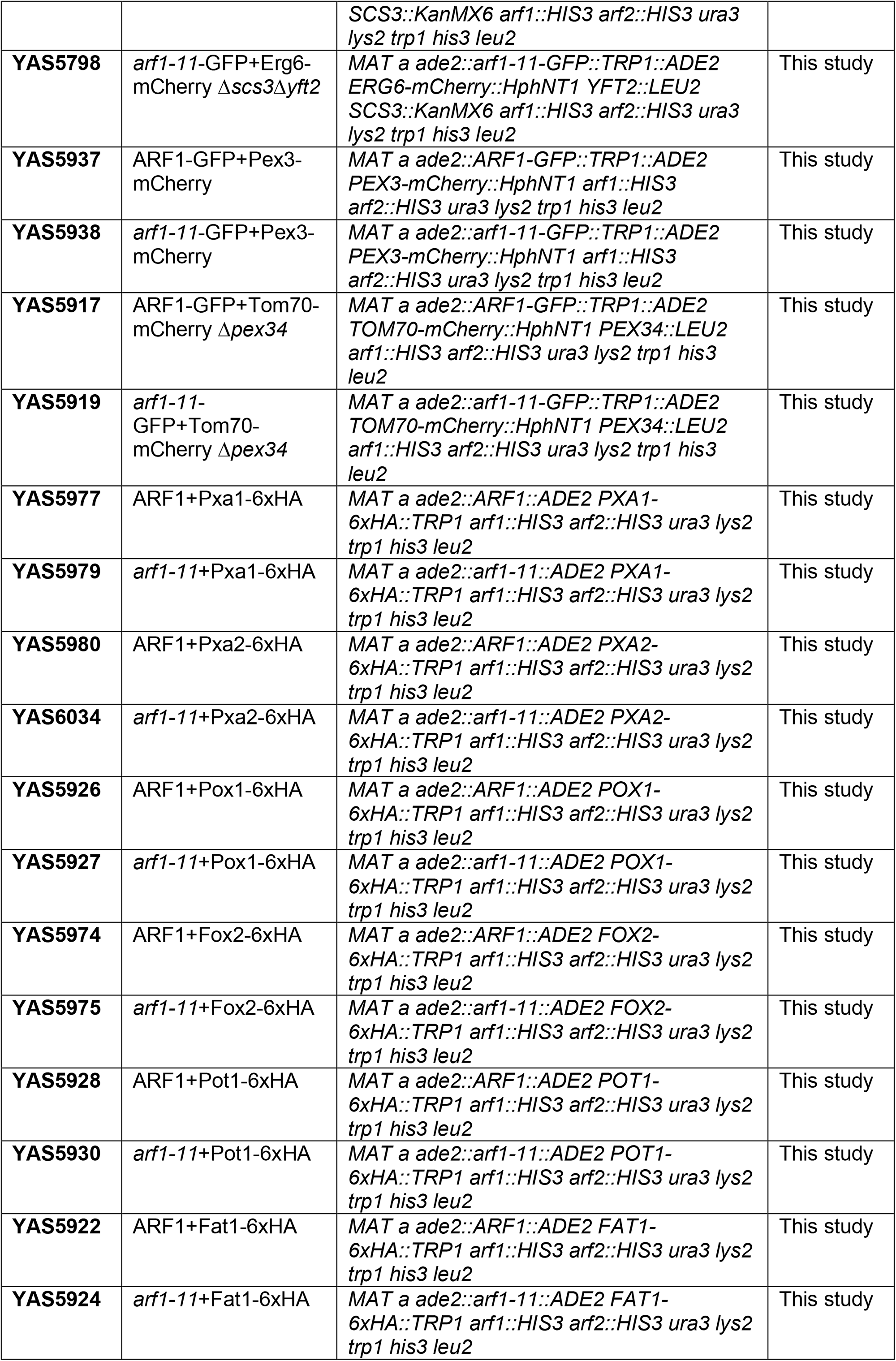
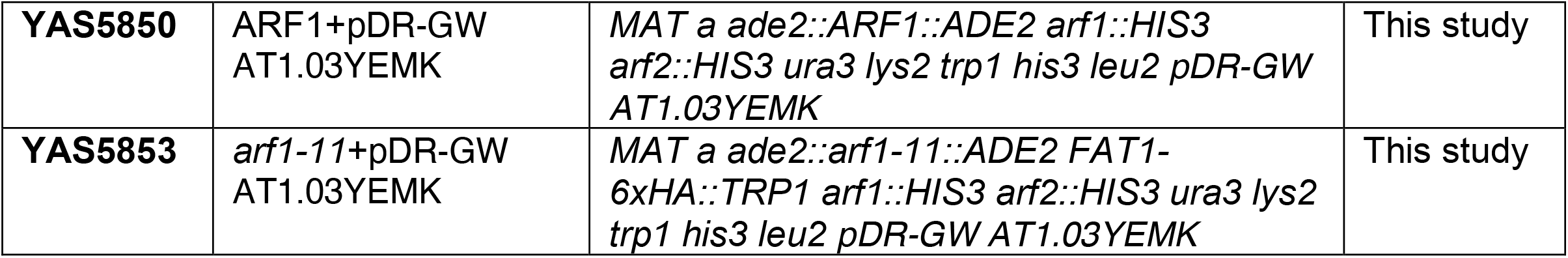
List of strains used in this study.

**Table 2.** List of plasmids used in this study.

| Construct | Insert | Promoter | Resistance/<br>Auxotrophy | Backbone | Copy<br>number | Source |
| --- | --- | --- | --- | --- | --- | --- |
| pGEX4-GGA2<sup>GAT</sup>-GST | Gga2<sup>GAT</sup> | Tac | Amp | pGEX4-T3 | - | Singer-Krüger., 2008 |
| pEGFP-N1 | - | CMV | Kan/Neo | pEGFP-N1 | - | This study |
| pEGFP-N1-Arf1 | Mammalian Arf1 (mArf1) | CMV | Kan/Neo | pEGFP-N1 | - | Addgene |
| pEGFP-N1-Arf1-11 | Mammalian *arf1-11* (mArf1-11) | CMV | Kan/Neo | pEGFP-N1 | - | This study |
| pGFP33-EV | - | - | Amp/URA3 | YCplac33 | CEN | This study |
| pGFP33-Arf1p-ARF1 | *ARF1* | *ARF1* | Amp/URA3 | YCplac33 | CEN | This study |
| pGFP33-Arf1p-ARF1<sup>Q71L</sup> | *ARF1* (Q71L) | *ARF1* | Amp/URA3 | YCplac33 | CEN | This study |
| pGFP33-Arf1p-ARF1<sup>T31N</sup> | *ARF1* (T31N) | *ARF1* | Amp/URA3 | YCplac33 | CEN | This study |
| pGFP33-Arf1p-ARF1<sup>K38T</sup> | *ARF1* (K38T) | *ARF1* | Amp/URA3 | YCplac33 | CEN | This study |
| pGFP33-Arf1p-ARF1<sup>L173S</sup> | *ARF1* (L173S) | *ARF1* | Amp/URA3 | YCplac33 | CEN | This study |
| pGFP33-Arf1p-ARF1<sup>K38T-E132D</sup> | *ARF1* (K38T+E132D) | *ARF1* | Amp/URA3 | YCplac33 | CEN | This study |
| pGFP33-Arf1p-ARF1<sup>K38T-E132D-L173S</sup> | *ARF1* (K38T+E132D+L173S) | *ARF1* | Amp/URA3 | YCplac33 | CEN | This study |
| ER-ΔN17-ARF1 | *SEC66-ARF1(ΔN17)* | *ARF1* | Amp/URA3 | YCplac33 | CEN | This study |
| ER-ΔN17-ARF1<sup>K38T-E132D-L173S</sup> | *SEC66-ARF1(ΔN17 ; (K38T+E132D+L173S))* | *ARF1* | Amp/URA3 | YCplac33 | CEN | This study |
| pDR-GW-AT1.03YEMK | *AT1.03YEMK+mVenus-mseCFP* | *PMA1* | Amp/URA3 | pDRf1GW-ura3 | 2μ | Addgene |

## References

1. Spang, A., Means of intracellular communication: touching, kissing, fusing. Microb Cell, 2021. 8(5): p. 87–90.

2. Zung, N. and M. Schuldiner, New horizons in mitochondrial contact site research. Biol Chem, 2020. 401(6-7): p. 793–809.

3. Silva, B.S.C., et al., Maintaining social contacts: The physiological relevance of organelle interactions. Biochim Biophys Acta Mol Cell Res, 2020. 1867(11): p. 118800.

4. Tamura, Y., S. Kawano, and T. Endo, Organelle contact zones as sites for lipid transfer. J Biochem, 2019. 165(2): p. 115–123.

5. Wu, H., P. Carvalho, and G.K. Voeltz, Here, there, and everywhere: The importance of ER membrane contact sites. Science, 2018. 361(6401).

6. Spang, A. and S. Mayor, Editorial Overview: Membranes and organelles: rethinking membrane structure, function and compartments. Curr Opin Cell Biol, 2018. 53: p. A1–A3.

7. Stefan, C.J., et al., Membrane dynamics and organelle biogenesis-lipid pipelines and vesicular carriers. BMC Biol, 2017. 15(1): p. 102.

8. Petkovic, M., C.E. O’Brien, and Y.N. Jan, Interorganelle communication, aging, and neurodegeneration. Genes Dev, 2021. 35(7-8): p. 449–469.

9. Cohen, Y., et al., Peroxisomes are juxtaposed to strategic sites on mitochondria. Mol Biosyst, 2014. 10(7): p. 1742–8.

10. Pu, J., et al., Interactomic study on interaction between lipid droplets and mitochondria. Protein Cell, 2011. 2(6): p. 487–96.

11. Schuldiner, M. and M. Bohnert, A different kind of love - lipid droplet contact sites. Biochim Biophys Acta Mol Cell Biol Lipids, 2017. 1862(10 Pt B): p. 1188–1196.

12. Shai, N., et al., Systematic mapping of contact sites reveals tethers and a function for the peroxisome-mitochondria contact. Nat Commun, 2018. 9(1): p. 1761.

13. Herms, A., et al., AMPK activation promotes lipid droplet dispersion on detyrosinated microtubules to increase mitochondrial fatty acid oxidation. Nat Commun, 2015. 6: p. 7176.

14. Rambold, A.S., S. Cohen, and J. Lippincott-Schwartz, Fatty acid trafficking in starved cells: regulation by lipid droplet lipolysis, autophagy, and mitochondrial fusion dynamics. Dev Cell, 2015. 32(6): p. 678–92.

15. Cui, L. and P. Liu, Two Types of Contact Between Lipid Droplets and Mitochondria. Front Cell Dev Biol, 2020. 8: p. 618322.

16. Chang, C.L., et al., Spastin tethers lipid droplets to peroxisomes and directs fatty acid trafficking through ESCRT-III. J Cell Biol, 2019. 218(8): p. 2583–2599.

17. Binns, D., et al., An intimate collaboration between peroxisomes and lipid bodies. J Cell Biol, 2006. 173(5): p. 719–31.

18. Kim, S., et al., Dysregulation of mitochondria-lysosome contacts by GBA1 dysfunction in dopaminergic neuronal models of Parkinson’s disease. Nat Commun, 2021. 12(1): p. 1807.

19. Dziurdzik, S.K. and E. Conibear, The Vps13 Family of Lipid Transporters and Its Role at Membrane Contact Sites. Int J Mol Sci, 2021. 22(6).

20. Spang, A., ARF1 regulatory factors and COPI vesicle formation. Curr Opin Cell Biol, 2002. 14(4): p. 423–7.

21. Arakel, E.C., et al., Dissection of GTPase-activating proteins reveals functional asymmetry in the COPI coat of budding yeast. J Cell Sci, 2019. 132(16).

22. Trautwein, M., et al., Arf1p provides an unexpected link between COPI vesicles and mRNA in Saccharomyces cerevisiae. Mol Biol Cell, 2004. 15(11): p. 5021–37.

23. Kilchert, C., et al., Defects in the secretory pathway and high Ca2+ induce multiple P-bodies. Mol Biol Cell, 2010. 21(15): p. 2624–38.

24. Jewell, J.L., et al., Metabolism. Differential regulation of mTORC1 by leucine and glutamine. Science, 2015. 347(6218): p. 194–8.

25. Meng, D., et al., Glutamine and asparagine activate mTORC1 independently of Rag GTPases. J Biol Chem, 2020. 295(10): p. 2890–2899.

26. Ackema, K.B., et al., The small GTPase Arf1 modulates mitochondrial morphology and function. EMBO J, 2014. 33(22): p. 2659–75.

27. Walch, L., et al., GBF1 and Arf1 interact with Miro and regulate mitochondrial positioning within cells. Sci Rep, 2018. 8(1): p. 17121.

28. Nagashima, S., et al., Golgi-derived PI(4)P-containing vesicles drive late steps of mitochondrial division. Science, 2020. 367(6484): p. 1366–1371.

29. Yahara, N., et al., Multiple roles of Arf1 GTPase in the yeast exocytic and endocytic pathways. Mol Biol Cell, 2001. 12(1): p. 221–38.

30. Nakamura, N., et al., ADRP is dissociated from lipid droplets by ARF1-dependent mechanism. Biochem Biophys Res Commun, 2004. 322(3): p. 957–65.

31. Bartz, R., et al., Dynamic activity of lipid droplets: protein phosphorylation and GTP-mediated protein translocation. J Proteome Res, 2007. 6(8): p. 3256–65.

32. Beller, M., et al., COPI complex is a regulator of lipid homeostasis. PLoS Biol, 2008. 6(11): p. e292.

33. Soni, K.G., et al., Coatomer-dependent protein delivery to lipid droplets. J Cell Sci, 2009. 122(Pt 11): p. 1834–41.

34. Ellong, E.N., et al., Interaction between the triglyceride lipase ATGL and the Arf1 activator GBF1. PLoS One, 2011. 6(7): p. e21889.

35. Bouvet, S., et al., Targeting of the Arf-GEF GBF1 to lipid droplets and Golgi membranes. J Cell Sci, 2013. 126(Pt 20): p. 4794–805.

36. Wilfling, F., et al., Arf1/COPI machinery acts directly on lipid droplets and enables their connection to the ER for protein targeting. Elife, 2014. 3: p. e01607.

37. Passreiter, M., et al., Peroxisome biogenesis: involvement of ARF and coatomer. J Cell Biol, 1998. 141(2): p. 373–83.

38. Lay, D., et al., Binding and functions of ADP-ribosylation factor on mammalian and yeast peroxisomes. J Biol Chem, 2005. 280(41): p. 34489–99.

39. Anthonio, E.A., et al., Small G proteins in peroxisome biogenesis: the potential involvement of ADP-ribosylation factor 6. BMC Cell Biol, 2009. 10: p. 58.

40. Yofe, I., et al., Pex35 is a regulator of peroxisome abundance. J Cell Sci, 2017. 130(4): p. 791–804.

41. Zhdankina, O., et al., Yeast GGA proteins interact with GTP-bound Arf and facilitate transport through the Golgi. Yeast, 2001. 18(1): p. 1–18.

42. Matto, M., et al., Role for ADP ribosylation factor 1 in the regulation of hepatitis C virus replication. J Virol, 2011. 85(2): p. 946–56.

43. Ohsaki, Y., et al., Inhibition of ADP-ribosylation suppresses aberrant accumulation of lipidated apolipoprotein B in the endoplasmic reticulum. FEBS Lett, 2013. 587(22): p. 3696–702.

44. Pennauer, M., et al., Shared and specific functions of Arfs 1-5 at the Golgi revealed by systematic knockouts. J Cell Biol, 2022. 221(1).

45. Veenhuis, M., et al., Proliferation of microbodies in Saccharomyces cerevisiae. Yeast, 1987. 3(2): p. 77–84.

46. Hiltunen, J.K., et al., The biochemistry of peroxisomal beta-oxidation in the yeast Saccharomyces cerevisiae. FEMS Microbiol Rev, 2003. 27(1): p. 35–64.

47. Moir, R.D., et al., SCS3 and YFT2 link transcription of phospholipid biosynthetic genes to ER stress and the UPR. PLoS Genet, 2012. 8(8): p. e1002890.

48. Yap, W.S., et al., The yeast FIT2 homologs are necessary to maintain cellular proteostasis and membrane lipid homeostasis. J Cell Sci, 2020. 133(21).

49. Omura, S., Cerulenin. Methods Enzymol, 1981. 72: p. 520–32.

50. Twig, G. and O.S. Shirihai, The interplay between mitochondrial dynamics and mitophagy. Antioxid Redox Signal, 2011. 14(10): p. 1939–51.

51. De Vos, K.J., et al., Mitochondrial function and actin regulate dynamin-related protein 1-dependent mitochondrial fission. Curr Biol, 2005. 15(7): p. 678–83.

52. Yu, F.Y., et al., Citrinin induces apoptosis in HL-60 cells via activation of the mitochondrial pathway. Toxicol Lett, 2006. 161(2): p. 143–51.

53. Legros, F., et al., Mitochondrial fusion in human cells is efficient, requires the inner membrane potential, and is mediated by mitofusins. Mol Biol Cell, 2002. 13(12): p. 4343–54.

54. Miyazono, Y., et al., Uncoupled mitochondria quickly shorten along their long axis to form indented spheroids, instead of rings, in a fission-independent manner. Sci Rep, 2018. 8(1): p. 350.

55. Bermejo, C., et al., Dynamic analysis of cytosolic glucose and ATP levels in yeast using optical sensors. Biochem J, 2010. 432(2): p. 399–406.

56. Just, W.W. and J. Peranen, Small GTPases in peroxisome dynamics. Biochim Biophys Acta, 2016. 1863(5): p. 1006–13.

57. Ferreira, R., et al., Metabolic engineering of Saccharomyces cerevisiae for overproduction of triacylglycerols. Metab Eng Commun, 2018. 6: p. 22–27.

58. Salo, V.T., et al., Seipin Facilitates Triglyceride Flow to Lipid Droplet and Counteracts Droplet Ripening via Endoplasmic Reticulum Contact. Dev Cell, 2019. 50(4): p. 478–493 e9.

59. Janke, C., et al., A versatile toolbox for PCR-based tagging of yeast genes: new fluorescent proteins, more markers and promoter substitution cassettes. Yeast, 2004. 21(11): p. 947–62.

60. Jimenez-Rojo, N., et al., Conserved Functions of Ether Lipids and Sphingolipids in the Early Secretory Pathway. Curr Biol, 2020. 30(19): p. 3775–3787 e7.

61. Persson, L.B., V.S. Ambati, and O. Brandman, Cellular Control of Viscosity Counters Changes in Temperature and Energy Availability. Cell, 2020. 183(6): p. 1572–1585 e16.

62. Guerin, B., P. Labbe, and M. Somlo, Preparation of yeast mitochondria (Saccharomyces cerevisiae) with good P/O and respiratory control ratios. Methods Enzymol, 1979. 55: p. 149–59.

63. Emaus, R.K., R. Grunwald, and J.J. Lemasters, Rhodamine 123 as a probe of transmembrane potential in isolated rat-liver mitochondria: spectral and metabolic properties. Biochim Biophys Acta, 1986. 850(3): p. 436–48.

64. Somlo, M., Induction and repression of mitochondrial ATPase in yeast. Eur J Biochem, 1968. 5(2): p. 276–84.

