## Supplemental Material for "The small GTPase Arf1 regulates ATP synthesis and mitochondria homeostasis by modulating fatty acid metabolism"

#### Figure S1

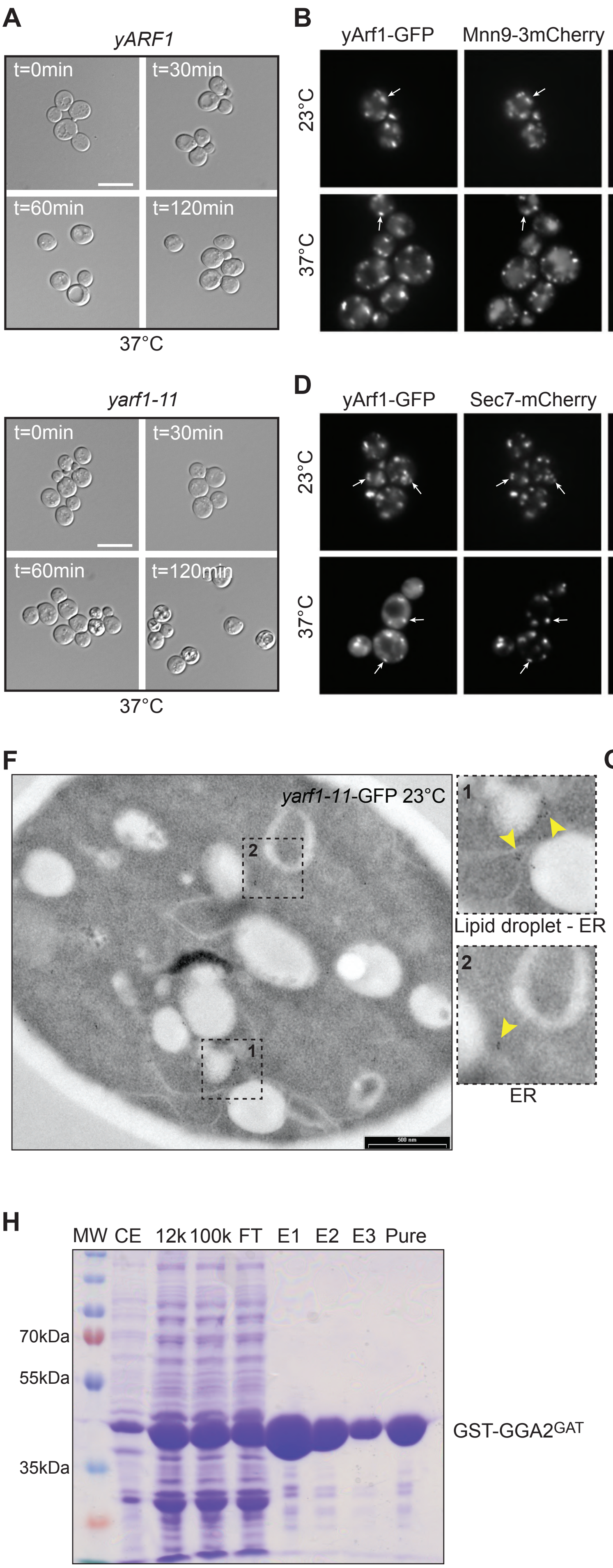
A
yARF1
t=0min
t=30min
t=60min
t=120min
37°C
yarf1-11
t=0min
t=30min
t=60min
t=120min
37°C
B
yArf1-GFP
Mnn9-3mCherry
Merge
23°C
37°C
C
yArf1-11-GFP
Mnn9-3mCherry
Merge
23°C
37°C
D
yArf1-GFP
Sec7-mCherry
Merge
23°C
37°C
E
yArf1-11-GFP
Tvp23-mCherry
Merge
23°C
37°C
F
yarf1-11-GFP 23°C
Lipid droplet - ER
ER
G
yarf1-11-GFP 37°C
ER
Lipid droplet
H
MW
CE
12k
100k
FT
E1
E2
E3
Pure
70kDa
55kDa
35kDa
GST-GGA2GAT

#### Figure S2

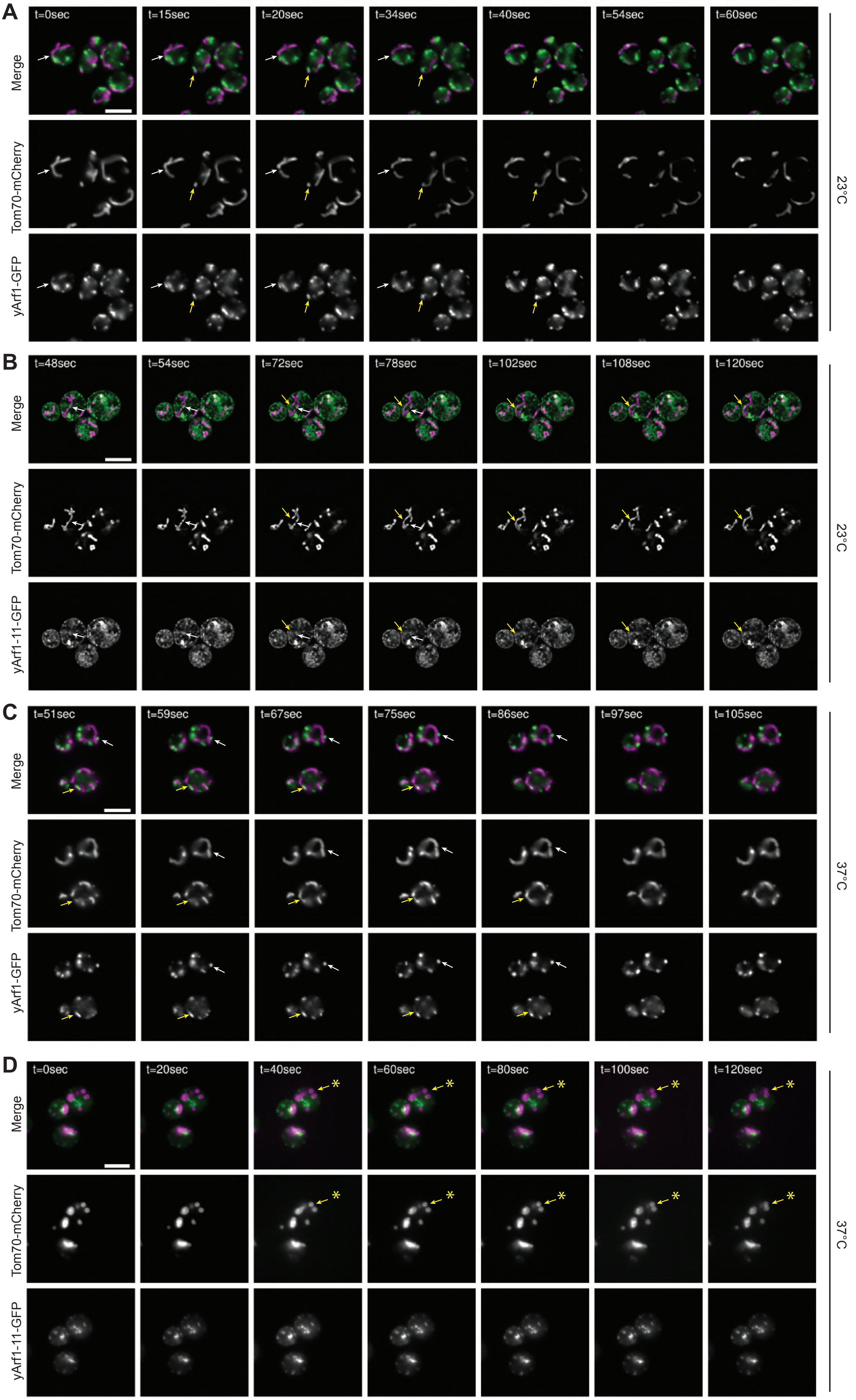

A
t=0sec
t=15sec
t=20sec
t=34sec
t=40sec
t=54sec
t=60sec
Merge
Tom70-mCherry
yArf1-GFP
23°C
B
t=48sec
t=54sec
t=72sec
t=78sec
t=102sec
t=108sec
t=120sec
Merge
Tom70-mCherry
yArf1-11-GFP
23°C
C
t=51sec
t=59sec
t=67sec
t=75sec
t=86sec
t=97sec
t=105sec
Merge
Tom70-mCherry
yArf1-GFP
37°C
D
t=0sec
t=20sec
t=40sec
t=60sec
t=80sec
t=100sec
t=120sec
Merge
Tom70-mCherry
yArf1-11-GFP
37°C

#### Figure S3

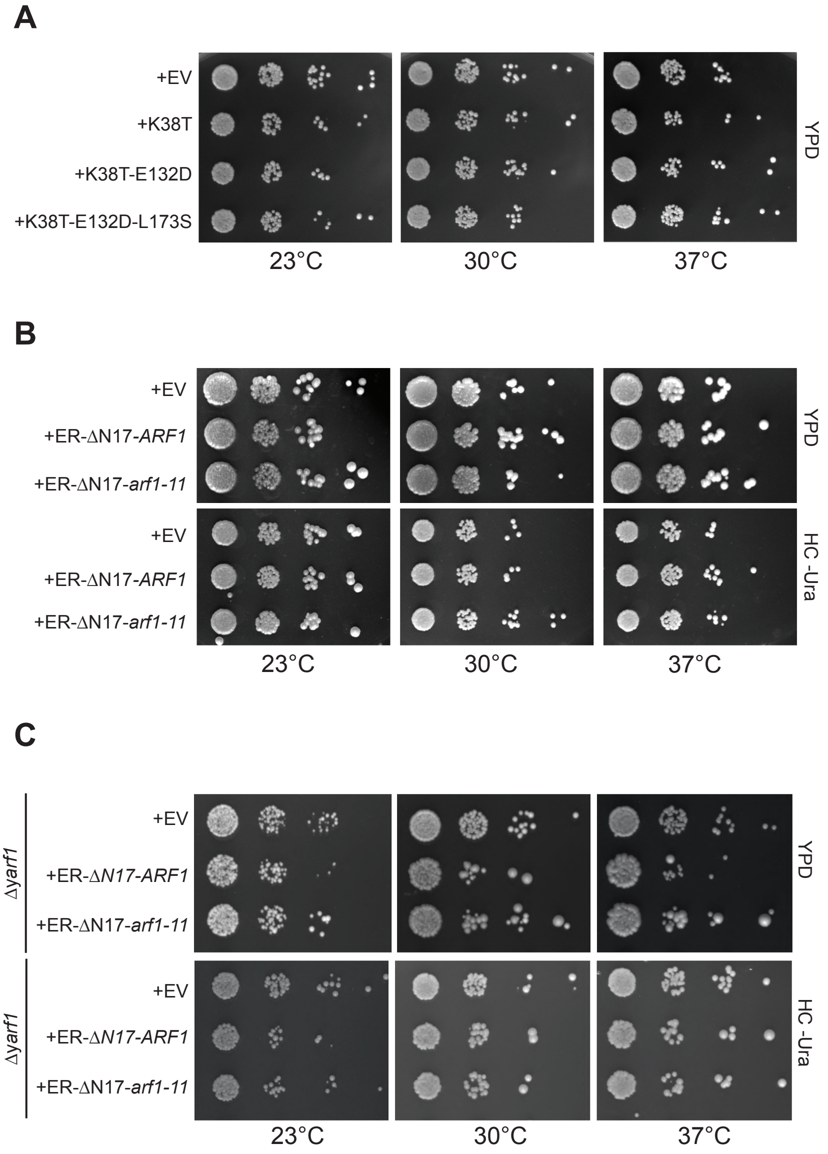

A
+EV
+K38T
+K38T-E132D
+K38T-E132D-L173S
YPD
23°C
30°C
37°C
B
+EV
+ER-ΔN17-ARF1
+ER-ΔN17-arf1-11
YPD
HC -Ura
23°C
30°C
37°C
C
Δyarf1
+EV
+ER-ΔN17-ARF1
+ER-ΔN17-arf1-11
YPD
HC -Ura
23°C
30°C
37°C

#### Figure S4

**A**

```
                          K38T
mArf1       MGNIFANLFKGLFGKKEMRILMVGLDAAGKTTILYKLKLGEIVTTIPTIGFNVETVEYKN 60
yArf1       MGLFASKLFSNLFGNKEMRILMVGLDGAGKTTVLYKLKLGEVITTIPTIGFNVETVQYKN 60
yarf1-11    MGLFASKLFSNLFGNKEMRILMVGLDGAGKTTVLYKLTLGEVITTIPTIGFNVETVQYKN 60
            ** : ::**..***:***********.*****:********::*************:***

mArf1       ISFTVWDVGGQDKIRPLWRHYFQNTQGLIFVVDSNDRERVNEAREELMRMLAEDELRDAV 120
yArf1       ISFTVWDVGGQDRIRSLWRHYYRNTEGVIFVVDSNDRSRIGEAREVMQRMLNEDELRNAA 120
yarf1-11    ISFTVWDVGGQDRIRSLWRHYYRNTEGVIFVVDSNDRSRIGEAREVMQRMLNEDELRNAA 120
            ************:**.*****::**:*:*********.*:.**** : *** *****:*.
                       E132D                                   L173S
mArf1       LLVFANKQDLPNAMNAAEITDKLGLHSLRHRNWYIQATCATSGDGLYEGLDWLSNQLRNQ 180
yArf1       WLVFANKQDLPEAMSAAEITEKLGLHSIRNRPWFIQATCATSGEGLYEGLEWLSNSLKNS 180
yarf1-11    WLVFANKQDLPDAMSAAEITEKLGLHSIRNRPWFIQATCATSGEGLYEGLEWSSNSLKNS 180
             **********:**.*****:******:*:* *:*********:******:****.*:*.

mArf1       K 181
yArf1       T 181
yarf1-11    T 181
            .
```

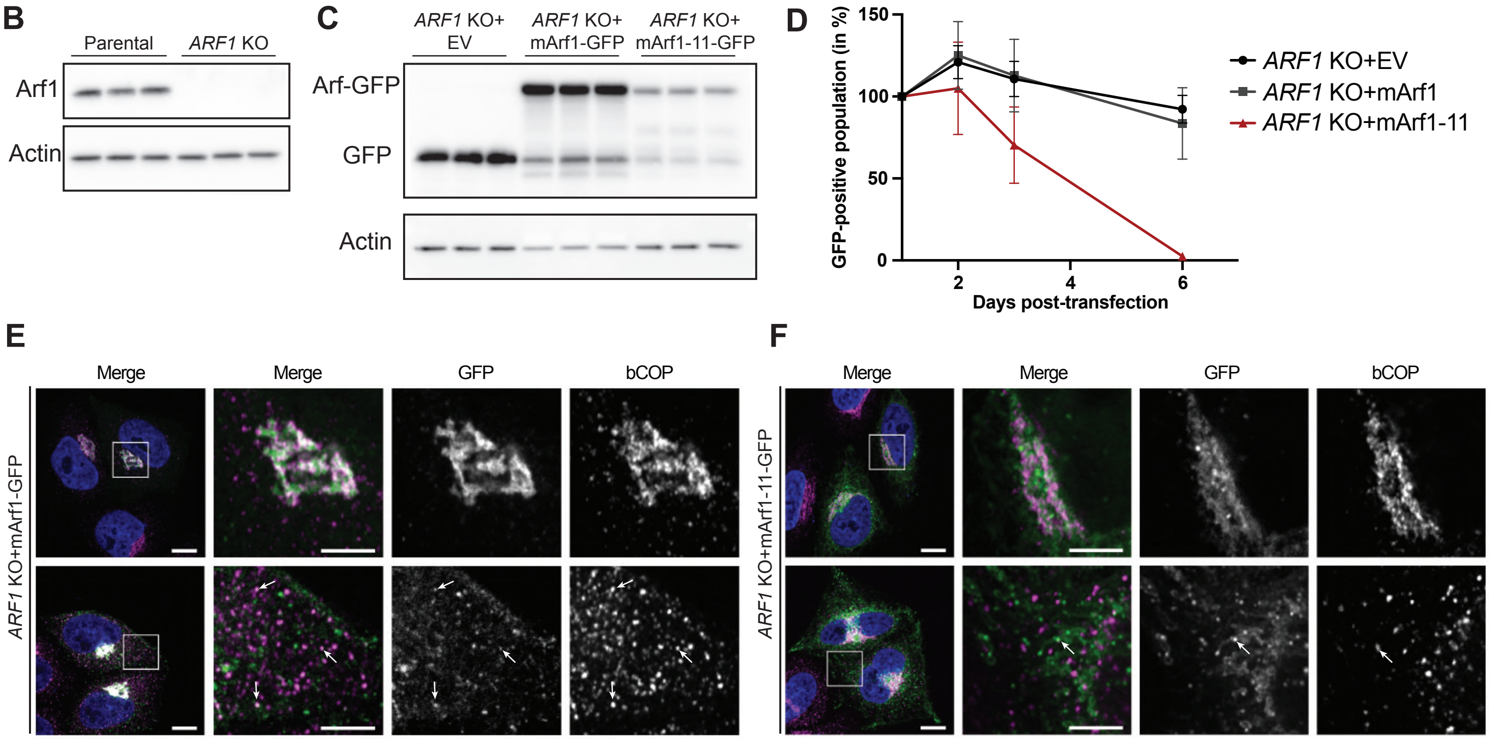

#### Figure S5

A

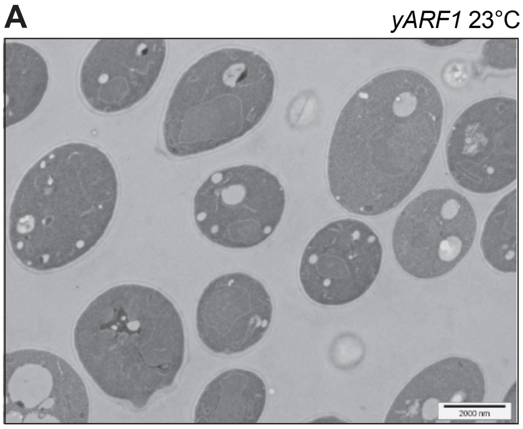
yARF1 23°C
2000 nm

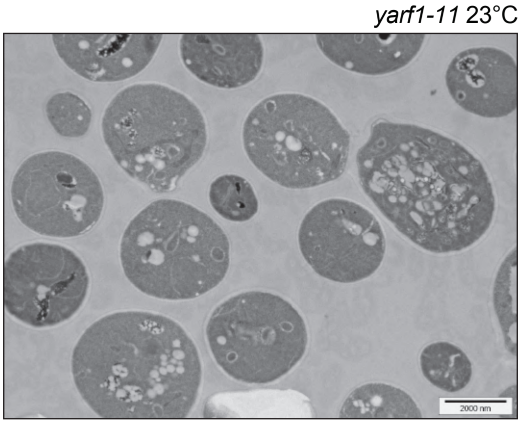
yarf1-11 23°C
2000 nm

B

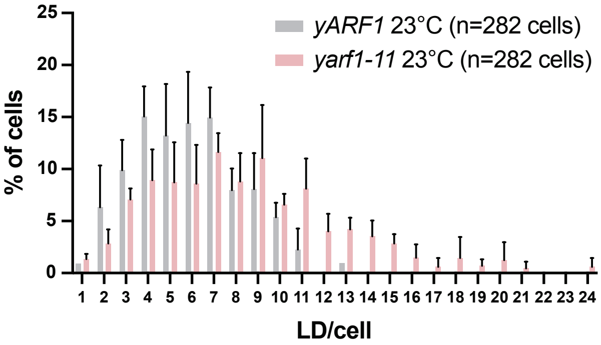
yARF1 23°C (n=282 cells)
yarf1-11 23°C (n=282 cells)
% of cells
LD/cell

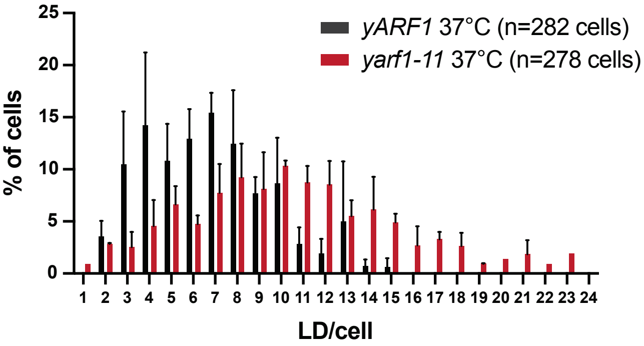
yARF1 37°C (n=282 cells)
yarf1-11 37°C (n=278 cells)
% of cells
LD/cell

### Figure S6

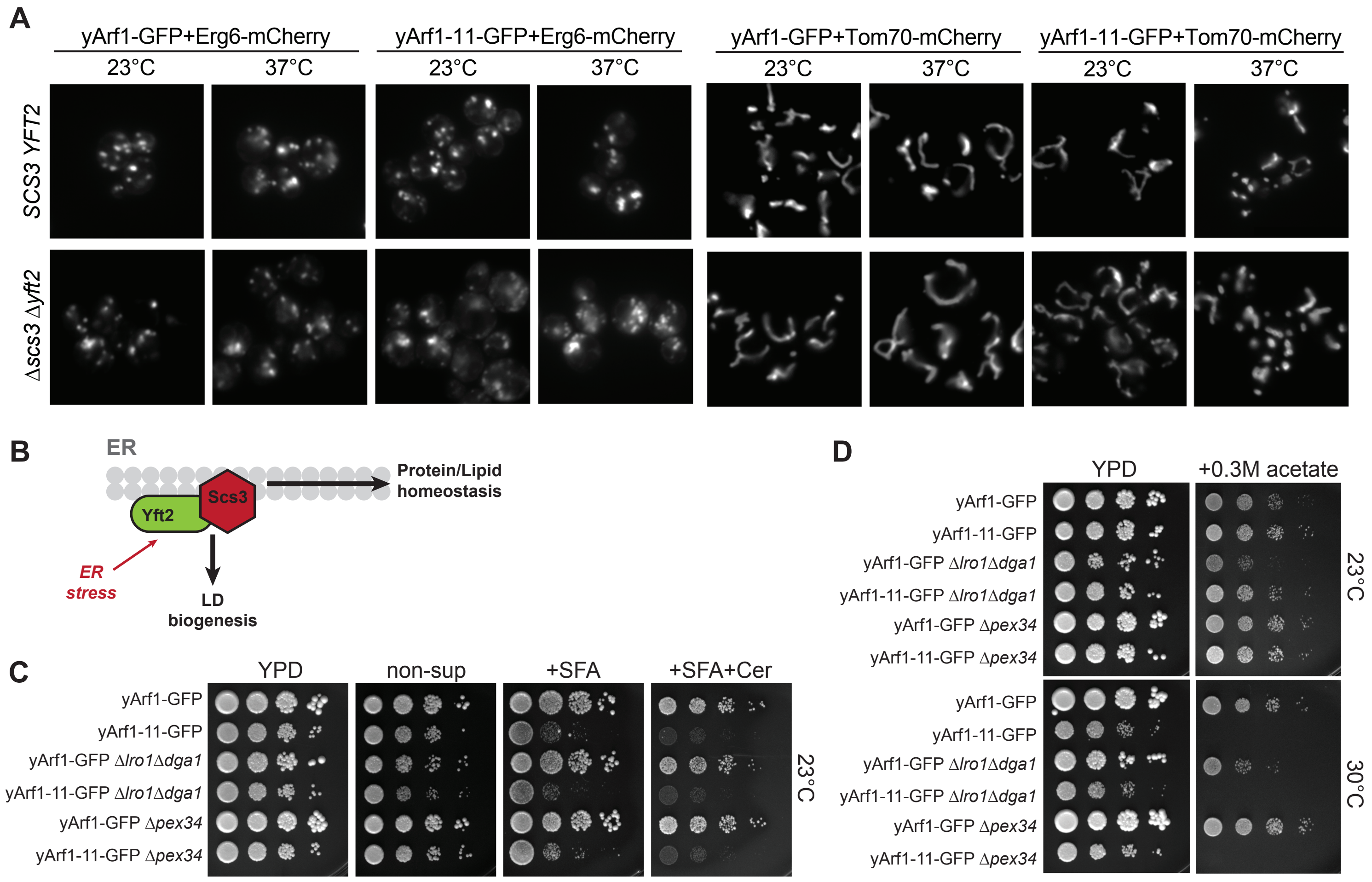
A
yArf1-GFP+Erg6-mCherry
yArf1-11-GFP+Erg6-mCherry
yArf1-GFP+Tom70-mCherry
yArf1-11-GFP+Tom70-mCherry
23°C
37°C
SCS3 YFT2
Δscs3 Δyft2
B
ER
Scs3
Yft2
Protein/Lipid homeostasis
ER stress
LD biogenesis
C
YPD
non-sup
+SFA
+SFA+Cer
yArf1-GFP
yArf1-11-GFP
yArf1-GFP Δlro1Δdga1
yArf1-11-GFP Δlro1Δdga1
yArf1-GFP Δpex34
yArf1-11-GFP Δpex34
23°C
D
YPD
+0.3M acetate
yArf1-GFP
yArf1-11-GFP
yArf1-GFP Δlro1Δdga1
yArf1-11-GFP Δlro1Δdga1
yArf1-GFP Δpex34
yArf1-11-GFP Δpex34
23°C
30°C

#### Figure S7

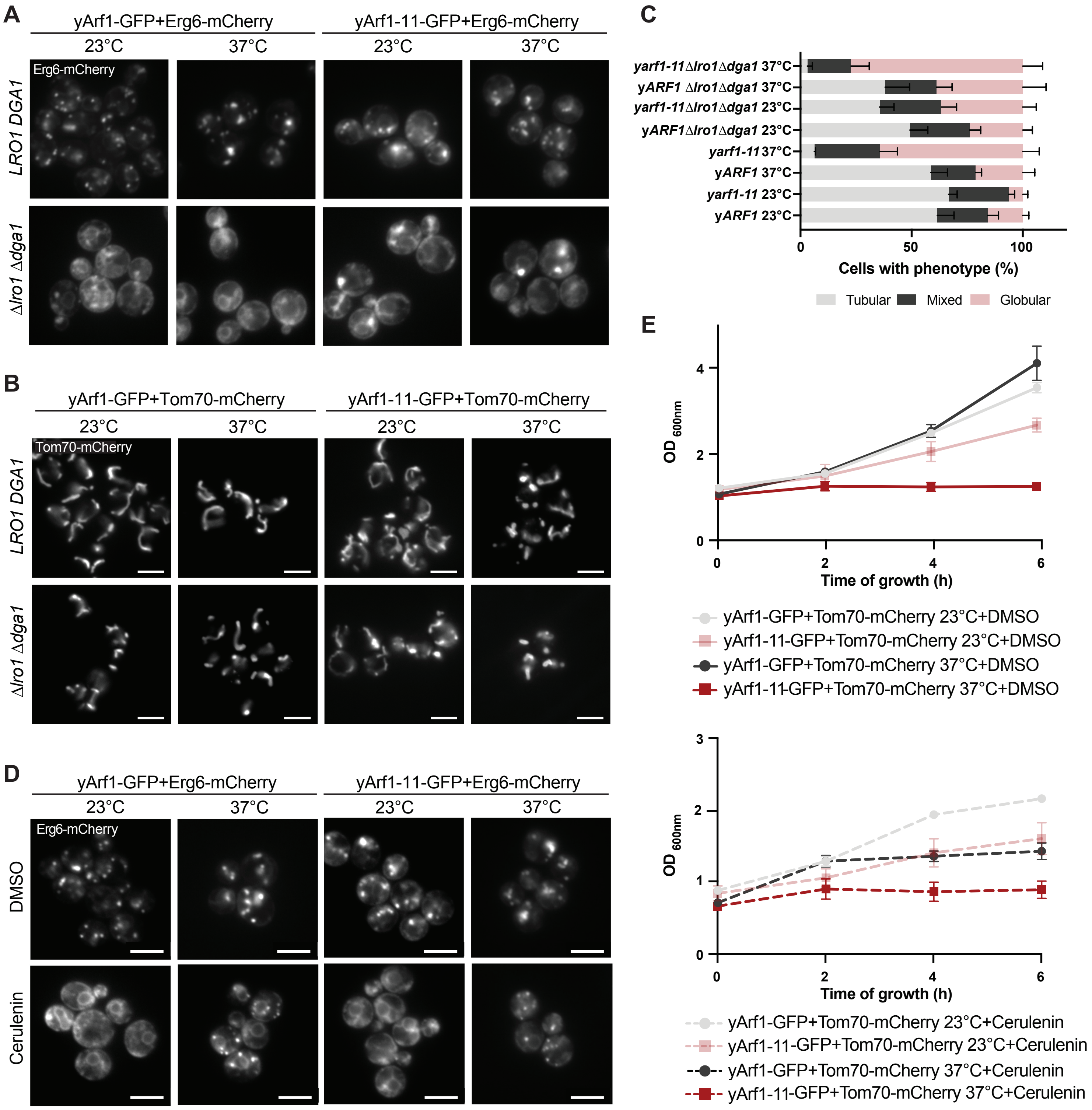

A
yArf1-GFP+Erg6-mCherry
yArf1-11-GFP+Erg6-mCherry
23°C
37°C
23°C
37°C
Erg6-mCherry
LRO1 DGA1
Δlro1 Δdga1
B
yArf1-GFP+Tom70-mCherry
yArf1-11-GFP+Tom70-mCherry
23°C
37°C
23°C
37°C
Tom70-mCherry
LRO1 DGA1
Δlro1 Δdga1
C
yarf1-11Δlro1Δdga1 37°C
yARF1 Δlro1Δdga1 37°C
yarf1-11Δlro1Δdga1 23°C
yARF1Δlro1Δdga1 23°C
yarf1-11 37°C
yARF1 37°C
yarf1-11 23°C
yARF1 23°C
0
50
100
Cells with phenotype (%)
Tubular
Mixed
Globular
D
yArf1-GFP+Erg6-mCherry
yArf1-11-GFP+Erg6-mCherry
23°C
37°C
23°C
37°C
Erg6-mCherry
DMSO
Cerulenin
E
OD 600nm
0
2
4
0
2
4
6
Time of growth (h)
yArf1-GFP+Tom70-mCherry 23°C+DMSO
yArf1-11-GFP+Tom70-mCherry 23°C+DMSO
yArf1-GFP+Tom70-mCherry 37°C+DMSO
yArf1-11-GFP+Tom70-mCherry 37°C+DMSO
OD 600nm
0
1
2
3
0
2
4
6
Time of growth (h)
yArf1-GFP+Tom70-mCherry 23°C+Cerulenin
yArf1-11-GFP+Tom70-mCherry 23°C+Cerulenin
yArf1-GFP+Tom70-mCherry 37°C+Cerulenin
yArf1-11-GFP+Tom70-mCherry 37°C+Cerulenin

### Figure S8

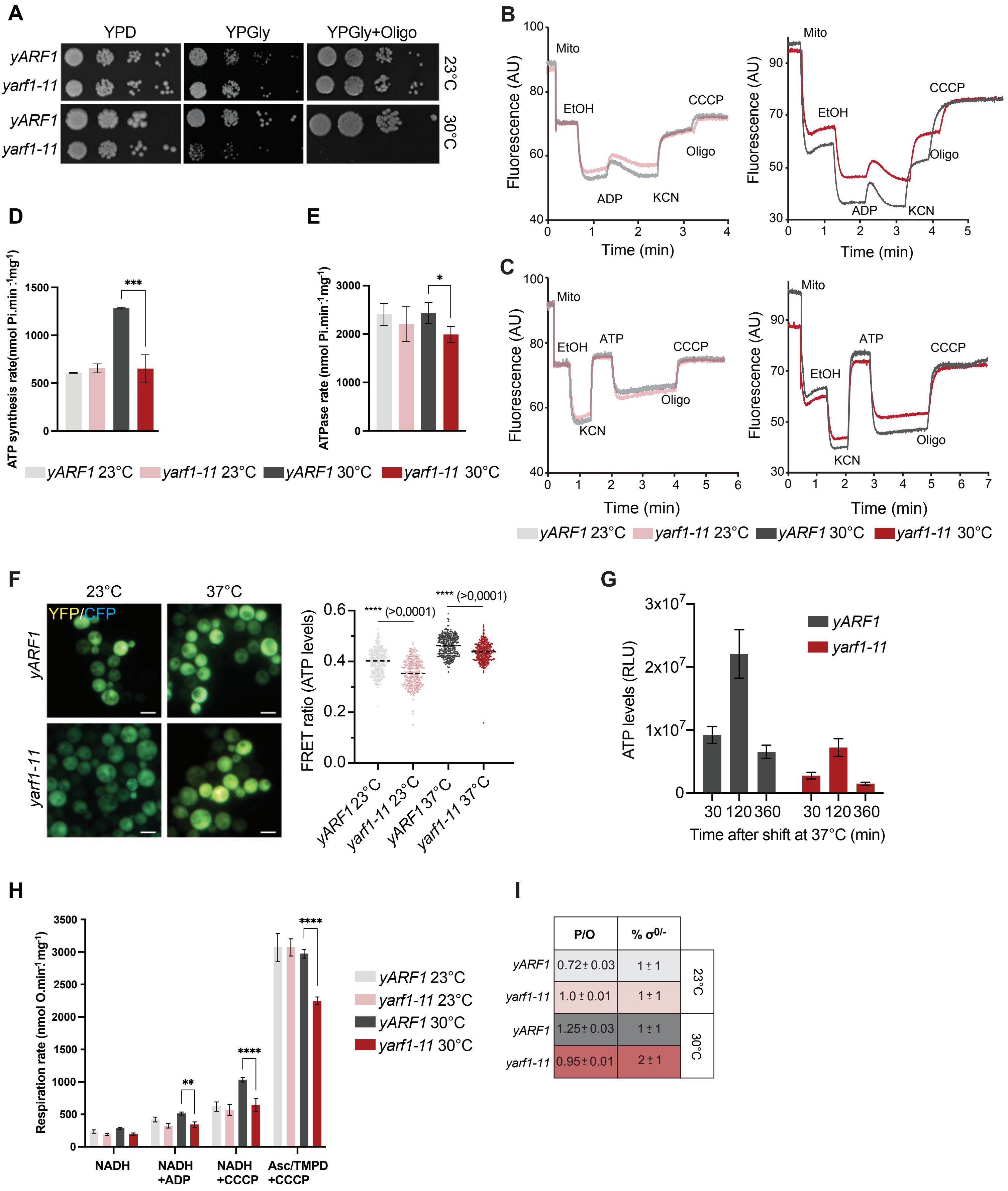


| | P/O | % $\sigma^{0/-}$ | |
|---|---|---|---|
| *yARF1* | 0.72 ± 0.03 | 1 ± 1 | 23°C |
| *yarf1-11* | 1.0 ± 0.01 | 1 ± 1 | |
| *yARF1* | 1.25 ± 0.03 | 1 ± 1 | 30°C |
| *yarf1-11* | 0.95 ± 0.01 | 2 ± 1 | |
